# The Mechanism of Dynamic Steady States in Lamellipodia

**DOI:** 10.1101/2024.11.18.624201

**Authors:** June Hyung Kim, Taeyoon Kim

**Author notes:** Correspondence should be addressed to Taeyoon Kim.

## Abstract

Lamellipodia are quasi-two-dimensional actin projections formed on the leading edge of the cell, playing an important role in sensing surrounding environments by forming focal adhesions. A branched actin network in the lamellipodia exhibits a stable, dynamic steady state characterized by a retrograde flow, which is attributed to a balance between network assembly at the leading edge and disassembly at the rear. Although the molecular players and architecture of the lamellipodia have been investigated extensively during recent decades, it still remains elusive how the dynamic steady state with continuous retrograde flow is achieved and robustly maintained. In this study, using an agent-based computational model, we probed how physical interactions between subcellular components in the lamellipodia lead to the dynamic steady state. We reproduced a steady retrograde flow induced by myosin activity and balance between network assembly and disassembly but hindered by resistances from adhesions formed on the underlying substrate. We demonstrated that different modes of dynamic steady states are possible, and that a network which failed to show the retrograde flow due to perturbations can be rescued by altering other factors. Our study provides insights into understanding how cells maintain the dynamic steady state of the lamellipodia in highly varying microenvironments.

## INTRODUCTION

Cell migration plays a central role in various biological processes, including morphogenesis, immune responses, wound repair, and cancer metastasis (1–3). One of the important steps for cell migration is cell protrusion, the outward extension of the plasma membrane. The cell protrusion allows cells to probe and sense surrounding environments by forming distinct actin architectures in response to different geometry of microenvironments. Two typical cell protrusions that emerge on cells migrating on a substrate are filopodia (finger-like) and lamellipodia (sheet-like). The lamellipodia comprise branched actin networks formed by Arp2/3 complex (4–8). The Arp2/3 complex is activated primarily via Wiskott–Aldrich syndrome protein (WASP) family, such as the verprolin-like central acidic (VCA) domain of the WAVE regulatory complex (9–12). After activation, Arp2/3 complex binds to the side of actin filaments (F-actins) to nucleate new filaments (branches) at the characteristic angle of ∼70° (13–16). Then, the branches elongate by polymerization with ATP-actin monomers. However, branch elongation does not last long since capping proteins quickly bind to elongating barbed ends to prevent further polymerization (17, 18). These elongating branches collectively push the cell membrane bearing tension outward, leading to the lamellipodial protrusion. Additionally, actin cross-linking proteins (ACPs), including α-actinin and filamin A, are known to help the branches push the membrane by physically interconnecting them (19, 20).

The branched network is also continuously pulled toward the rear by contractile forces generated by myosin motors. The network is accumulated into bundles called actin arcs found at the interface between lamellipodia and lamella (21, 22). F-actins undergo ATP hydrolysis during movement toward the rear, becoming ADP-F-actin that is more prone to depolymerization and severing induced by cofilin which is also known as Actin Depolymerization Factor (ADF) (23–25). As a result of F-actin disassembly, the actin arc is able to maintain a relatively consistent thickness rather than thickening over time from continuous accumulation of F-actins (22). Actin monomers generated from F-actin disassembly bind to profilin to avoid de novo (spontaneous) nucleation and switch to ATP-G-actin more rapidly (26–28). These profilin-bound actin monomers are further transported to the leading edge to be used for the elongation of new branches. Coordinated, balanced activities of these proteins result in a dynamic steady state characterized by the actin retrograde flow from the leading edge to the rear. F-actins in the lamellipodia keep forming focal adhesions (FAs) on an underlying substrate via integrin proteins, which enables cells to exert traction forces required for mechanosensing and cell migration and to impede the actin retrograde flow (29–32).

The dynamic steady state of branched actin networks in the lamellipodia has been understood relatively well. Nevertheless, none of the reconstituted systems consisting of purified cytoskeletal proteins have succeeded in reproducing the dynamic steady state. Systems with cell extract encapsulated in water-in-oil droplets showed persistent contractile flows (33), but the exact contribution of each protein in the cell extract to mediating the flow and maintaining its dynamic steady state is still unclear. Thus, the exact recipe of proteins required for the dynamic steady state observed in the lamellipodia is still elusive. In addition, this dynamic steady state is surprisingly robust. Lamellipodia constantly adapt to changes in environmental cues (e.g., stiffness and adhesion conditions) by varying their dynamic steady state (34, 35). Despite experimental observations, how the lamellipodia show such high adaptability to environments remains to be explained.

Computational models can help illuminate the mechanisms of the dynamic steady state exhibited by branched actin networks in the lamellipodia and their adaptability by overcoming intrinsic limitations of experiments. Indeed, there have been several mathematical and computational models designed to describe the biochemical and mechanical dynamics of the branched actin network in the lamellipodia with consideration of actin turnover, protrusive forces, motor activity, FA dynamics, and different types of substrates (36–44). A recent discrete model showed how the retrograde flow speed and morphology of branched actin networks are affected by physical interactions with a single matured FA site (45). We recently developed a filament-level model to reproduce the actin retrograde flow induced by myosin activity against resistances from many nascent FA sites formed on an underlying substrate, but the flow took place only once without a steady state due to the absence of actin dynamics (18). Despite these previous efforts, there has not been any model capable of reproducing the dynamic steady state of lamellipodia without drastic simplification of cytoskeletal elements and structures.

In this study, we developed a rigorous model for the lamellipodia with detailed descriptions of physical interactions between key molecular players identified in experiments. Our model was able to reproduce the steady-state retrograde flow emerging from a balance between network assembly at the leading edge, network disassembly and motor activity at the rear, and nascent FAs forming between the network and the underlying substrate. However, when this balance was perturbed by a change in one of factors, the dynamic steady state disappeared. We further demonstrated how the dynamic steady state can be achieved again to adapt to the perturbation. Our study provides insights into understanding the resilient maintenance and recovery of the dynamic steady state observed in the lamellipodia.

## MATERIALS AND METHODS

An agent-based model is developed to simulate dynamic interactions between key molecular players in the lamellipodia, including F-actin, molecular motor, Arp2/3 complex, and ACP (Fig. 1A). In our model, F-actin is simulated as serially connected cylindrical segments with polarity defined by the barbed and pointed ends. Arp2/3 and ACP consist of two segments connected in series, whereas motors are simplified into a backbone connected to 8 motor arms. Each of the motor arms kinetically represents 4 myosin heads. F-actin can undergo de novo nucleation, polymerization, depolymerization, and angle-dependent severing. Arp2/3 binds to the side of existing F-actin to nucleate a new filament at the characteristic branching angle of 70°. ACP connects pairs of F-actins to form functional cross-linking points. Each motor arm binds to F-actin and walks toward the barbed end of F-actin, and it also unbinds from F-actin in a force-dependent manner. An underlying substrate is coarse-grained into a triangulated mesh with nodes whose z positions are fixed at z = 0 (Fig. 1B). Nascent FAs are mimicked by transiently forming elastic links between the endpoints of actin segments and the substrate nodes (Fig. 1B). The displacements of substrate nodes and all the segments constituting F-actin, Arp2/3, ACP, and motor are determined by the Langevin equation based on Brownian dynamics. To mimic the geometry of the lamellipodia, a relatively thin 3D domain (5×2.5×0.1 µm) is employed (Fig. 1C). The +y boundary and -y boundary are considered the leading edge and the interface between lamellipodia and lamella, respectively. At the beginning of simulation, a branched actin network is assembled from filament seeds formed on the -y boundary via de novo nucleation. ∼90% of actin is used for assembling the network. During network assembly, dynamic behaviors of F-actin, Arp2/3, ACP, motor, and FA described earlier take place except motor walking. Motors are located only near the -y boundary, and FAs are allowed to form within a specific region, unlike the other elements (Fig. 1C). After network assembly, motors start walking along F-actins, leading to a retrograde flow. In addition, F-actins are polymerized from the barbed ends only near the leading edge and depolymerized from the pointed ends only near the -y boundary (Fig. 1C). F-actins can also be severed anywhere at a rate proportional to a local bending angle. Even without any spatial constraint, F-actin severing predominantly takes place in the region where the network is compacted into a bundle (mimicking the actin arc) due to the motor activity. Free actin segments generated from F-actin disassembly and severing can be recycled for network assembly occurring near the leading edge, allowing for a continuous actin retrograde flow. All of these dynamic behaviors and interactions can allow the branched actin network to reach a dynamic steady state under specific conditions as shown later.

**Figure 1.**
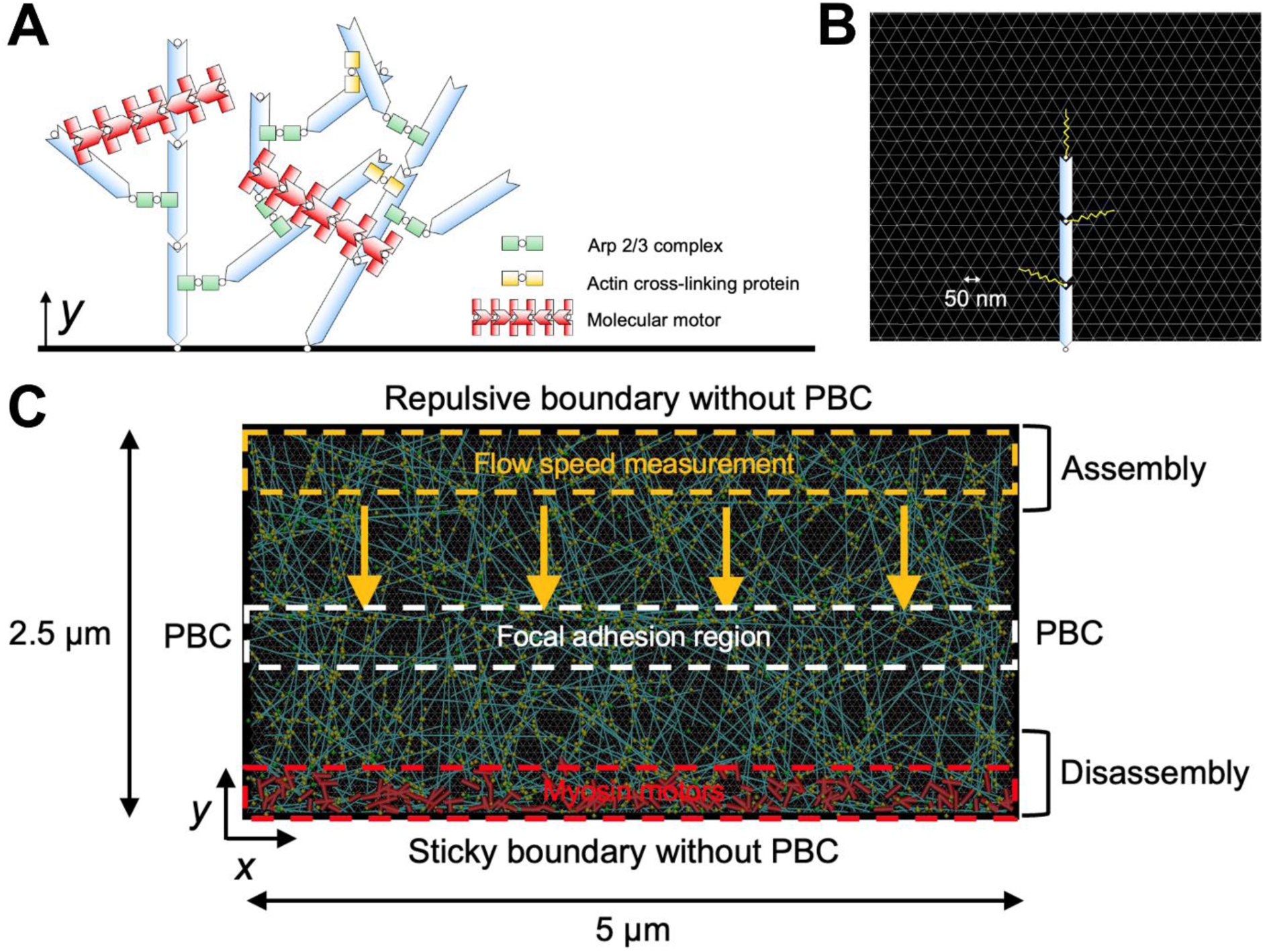
The lamellipodia model. (A) A branched network is created by self-assembly and interactions of F-actin (cyan), Arp2/3 complex (green), ACP (yellow), and motor (red), all of which are simplified via cylindrical segments. (B) An underlying substrate is simplified into a triangulated mesh with the chain length of 50 nm (gray). The endpoints of actin segments can transiently form an elastic link (yellow) to the substrate. (C) Description of different regions for dynamic events in the computational domain (5 µm×2.5 µm). Actin polymerization occurs only in the assembly region (y = 2.125 - 2.5 µm), whereas actin depolymerization takes place only in the disassembly region (y = 0 - 0.375 µm). Motors are initially located near the -y boundary (y = 0 - 0.1875 µm). The links between the substrate and the network can form only in the focal adhesion region (y = 0.325 - 0.675 µm). We use F-actins located near the +y boundary for the calculation of retrograde flow speed. The periodic boundary condition is applied in the x direction, whereas repulsive and sticky boundary conditions are applied to the +y and -y boundaries, respectively.

## RESULTS

Using our model, we explored a wide parametric space to find conditions for the continuous retrograde flow in the dynamic steady state. We identified a condition where the dynamic steady state was maintained over a long time. Under this reference condition, network morphology was similar regardless of time points (Fig. 2A). The number of FAs forming on the underlying substrate and the total force acting on the underlying substrate did not vary significantly over time (Figs. 2B-D). The retrograde flow speed remained steady spatiotemporally (Figs. 2E-G). In the following results, we varied the values of key parameters to understand how the dynamic steady state is altered and maintained.

**Figure 2.**
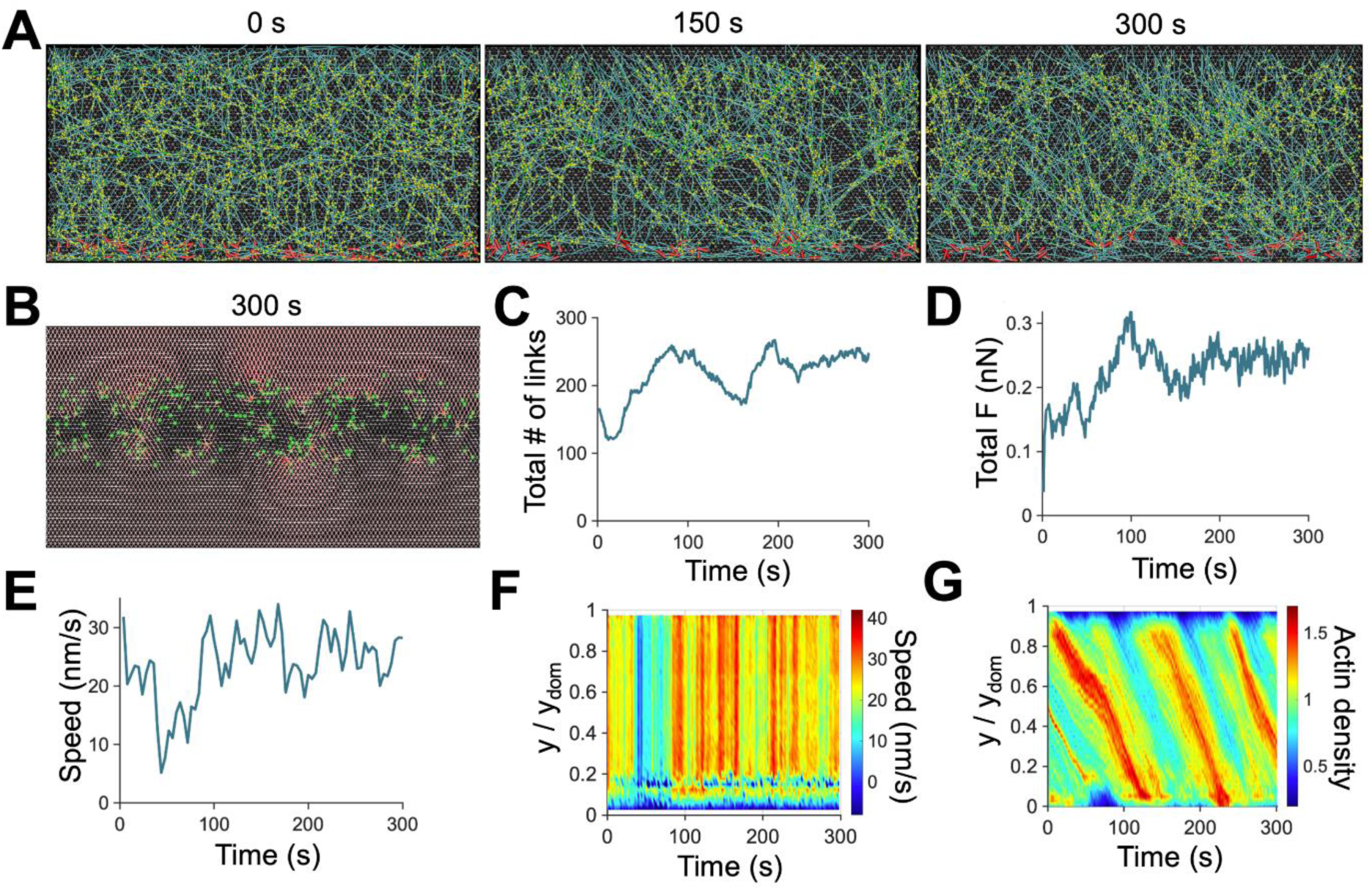
The dynamic steady state of the branched network under the reference condition. (A) Snapshots of networks taken at 0 s, 150 s, and 300 s. The network was homogeneous and maintained continuity regardless of time points. (B) Snapshot showing links (green circles) formed in the focal adhesion region (y = 0.81 - 1.69 µm) and forces acting on the underlying substrate via color scaling (red: high force, white: low force) taken at 300 s. (C) Time evolution of the number of the links formed between the substrate and the network. (D) The total substrate force exerted by the network via the links as a function of time. (E) Time evolution of retrograde flow speed. (F) Kymograph of flow speed as a function of y position and time. Quantities shown in (C-F) changes more at *t* < 100 s but exhibit smaller fluctuations around average values. (G) Kymograph of actin density as a function of y position and time. Under this reference condition, key parameter values are *k*_+,A_ = 12 µM^-1^s^-1^, *R*_ACP_ = 0.04, *R*_Arp2/3_ = 0.01, *k*_-,A_ = 6 s^-1^, *k*_0,sev_ = 10^-45^ s^-1^, *λ*_sev_ = 1.0 deg, *R*_M_ = 0.004, *k*_20_ = 17 s^-1^, and *A*_FA_ = 0.35.

### Proper assembly and connectivity of networks are required for maintaining the dynamic steady state

We first investigated the effects of a variation in parameters governing network assembly on the dynamic steady state. We found that actin polymerization occurring near the leading edge should be fast enough for maintaining a continuous branched network structure. When F-actins were polymerized too slow due to low actin polymerization rate constant (*k*_+,A_), the contracting network lost connection to newly assembled F-actins near the leading edge, resulting in the loss of a steady state (Figs. S2A, F, G). Slow actin polymerization further led to a pronounced oscillation in actin density (Figs. S2C, E). With *k*_+,A_ higher than 1 µM^-1^s^-1^, the continuity of the network structure was sustained in the flow direction, and the oscillation did not appear. The retrograde flow speed was also relatively steady only when the network showed continuity in the flow direction (Figs. S2F, G). A further increase in *k*_+,A_ did not make a difference in network behaviors because it was assumed in the model that the concentration of free actin monomers is enforced to stay above 10% of the total actin concentration (Figs. S2B, D-G). In other words, the availability of actin monomers provided from depolymerization and severing at the rear becomes a limiting factor for actin turnover if actin polymerization is sufficiently fast.

The influence of a variation in Arp2/3 density (*R*_Arp2/3_) was also tested because Arp2/3 plays a critical role in network assembly by forming new branches from existing F-actins near the leading edge. The lack of Arp2/3 caused a decrease in the flow speed since the formation of new branches was unable to keep up with the network disassembly occurring at the rear near the -y boundary (Figs. S3A, F, G). This led to the accumulation of F-actin and significant oscillation in actin density (Figs. S3C, E). High *R*_Arp2/3_ resulted in the formation of more branches on a fraction of vertically growing structures, so the network became more heterogeneous in the x direction with lower connectivity (Fig. S3B). Despite high heterogeneity in the x direction, *R*_Arp2/3_ above threshold level (*R*_Arp2/3_ ≥ 0.008) resulted in a continuous retrograde flow, and there was no significant effect of a further increase in *R*_Arp2/3_ on flow speed because network disassembly is the limiting factor as described earlier (Figs. S3D-G).

In our model, branches always form toward the leading edge. Thus, one filament seed created on the -y boundary grows as a relatively narrow (in the x direction) branched network up to the leading edge. Although all F-actins within this structure have superior connectivity via permanently bound Arp2/3, connection between structures originating from different filament seeds does not exist. ACPs connect pairs of branches to enhance network connectivity in the x direction. To understand the importance of network connectivity in the direction perpendicular to the flow, we varied ACP density (*R*_ACP_). With sufficiently high ACP density (*R*_ACP_ ≥ 0.02), the network could contract as a single entity and maintain a steady retrograde flow (Figs. S4B, D-G). With low ACP density (*R*_ACP_ < 0.02), forces generated by motors were not uniformly transmitted through an entire network due to lack of network connectivity (Fig. S4A). Thus, the network could not contract as a whole and was stalled due to frictional forces from FAs (Figs. S4C, E-G). In sum, the sufficiently fast assembly of a well-connected network is required for the dynamic steady state with a continuous retrograde flow and uniform network contraction.

### Network disassembly via depolymerization and severing is critical for maintaining the dynamic steady state

Considering the importance of actin disassembly as the limiting factor for actin turnover, we next probed how network morphology and retrograde flow are regulated by the depolymerization and severing of F-actins. First, the depolymerization rate was varied by changing the value of actin depolymerization rate (*k*_-,A_). We found that the dynamic steady state with physiologically relevant retrograde flow speed emerged when the actin depolymerization was neither too fast nor too slow. The absence of actin disassembly (*k*_-,A_ = 0 s^-1^) resulted in the accumulation of F-actins towards the -y boundary, corresponding to one-time, irreversible contraction of the branched network (Figs. 3A, C), which we showed in our recent study (18). The flow speed decayed over time as F-actin accumulated towards the -y boundary (Figs. 3F and S5A). By contrast, cases with very fast depolymerization (*k*_-,A_ ≥ 8 s^-1^) led to the rapid depolymerization of F-actins prior to motor-mediated network contraction, causing disconnection between the network and the motors (Figs. 3B, D). As a result, the retrograde flow disappeared after 100 s (Figs. 3F and S5A). With intermediate depolymerization rates (*k*_-,A_ = 4-6 s^-1^), the flow speed was high and steady, and the network showed relatively homogeneous actin distribution similar to the reference case (Figs. 3E, F).

**Figure 3.**
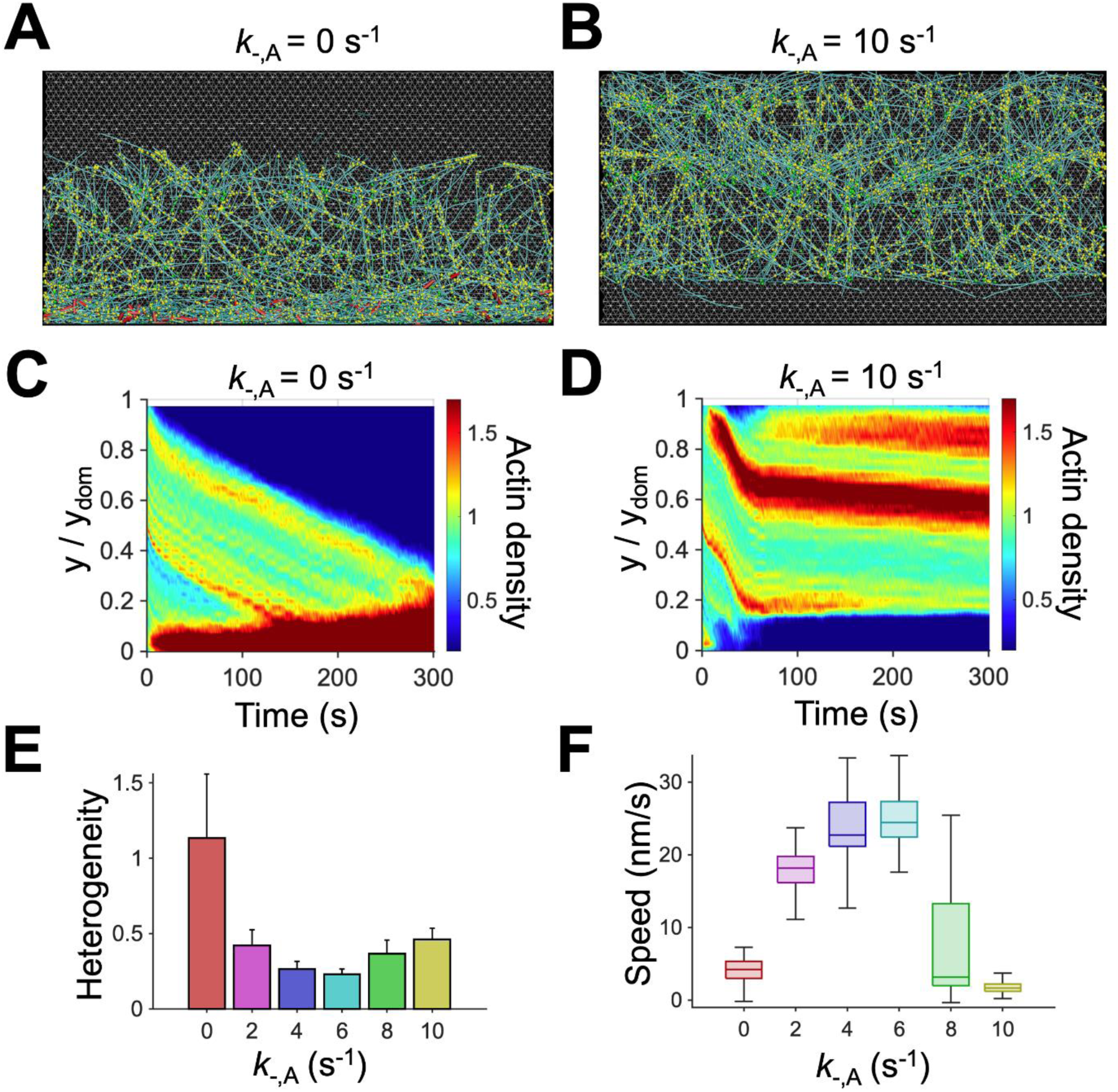
Intermediate actin depolymerization rate led to a continuous flow with uniform network morphology. (A, B) Snapshots of branched networks taken at ∼150 s with a lower or higher value of actin depolymerization rate (*k*_-,A_) relative to that of the reference condition, 6 s^-1^. Network contraction without actin depolymerization led to loss of connection between the network and the leading edge, whereas too fast actin depolymerization resulted in loss of connection between the network and the motors. (C, D) Kymographs of actin density as a function of y position and time with different *k*_-,A_. (E) Heterogeneity of the network quantified as a coefficient of variation in actin density in the y direction. A lower value is indicative of a more homogeneous network. (F) Retrograde flow speed for different values of *k*_-,A_. With intermediate values of *k*_-,A_, the network was more homogenous, and the flow speed was comparable with experimental observations.

It was shown experimentally that F-actin could be severed at an angle-dependent rate during thermal fluctuation (46). In our previous study, we have shown that angle-dependent F-actin severing can drive the reversible (i.e., pulsatile) contraction of actomyosin networks by enhancing the disassembly of aggregating networks (47). In our model, it is assumed that F-actin severing occurs in an angle-dependent manner in an entire domain. As before, the severing rate is determined by two parameters: actin severing rate constant (*k*_0,sev_) and angle sensitivity (*λ*_sev_) (Eq. S9). We probed the effects of a variation in *k*_0,sev_ and *λ*_sev_ on the dynamic steady state. The dynamic steady state was maintained well with steady flow speed and homogeneous network morphology only when *k*_0,sev_ was at intermediate level, *k*_0,sev_ = 10^-55^–10^-45^ (Figs. 4E, F and S5B). The absence of severing (*k*_0,sev_ = 0) led to the stalling of network contraction at early times (Figs. 4A, C). As F-actins in the motor region were disassembled, the network eventually lost connection to the motors. As a result, the branched network failed to reach a steady state and became noticeably heterogenous with a high variation in F-actin density in the y direction (Figs. 4C, E, F). Highly branched, cross-linked networks are hard to contract into bundles due to their high compressive resistance; the length of buckling units in these networks is very short, so large contractile forces are necessary to buckle them for network contraction. The angle-dependent severing can make the network contraction easier. Thus, without severing, the network contraction was stalled. Frequent F-actin severing (*k*_0,sev_ = 10^-25^) induced rapid fragmentation of F-actins in the motor region, where F-actins experience an increase in their bending angles due to buckling induced by network contraction (Figs. 4B, D, F). This led to rapid loss of connection between the network and the motors. Thus, the flow speed and the actin density showed biphasic dependence on *k*_0,sev_.

**Figure 4.**
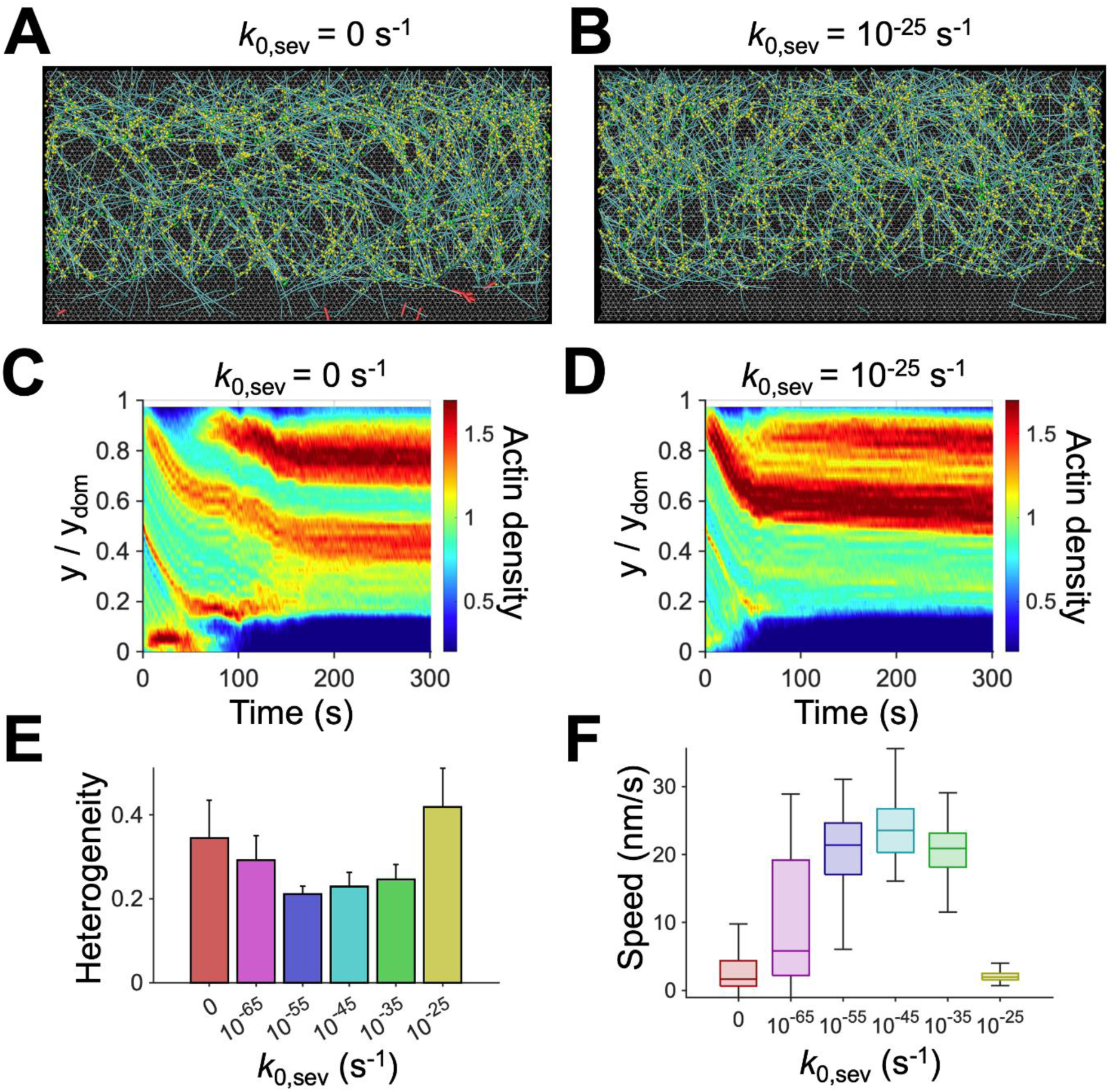
Intermediate F-actin severing activity is necessary to maintain a dynamic steady state. (A, B) Snapshots of the branched networks taken at ∼150 s with a lower or higher value of severing rate constant (*k*_0,sev_) relative to that of the reference condition, 10^-45^ s^-1^. In the absence of severing, a bundle structure was formed by the initial accumulation of F-actins. Due to the delayed disassembly of the bundle, motors remained trapped within the bundle until ∼60 s and could not induce further network contraction. Once the bundle was finally disassembled by depolymerization, motors were unable to find F-actins to bind, leading to disconnection between the network and the motors. With fast severing activity, the filaments were immediately disassembled before motors could induce contraction, and the network eventually lost connection to the motors. (C, D) Kymographs of actin density as a function of y position and time with different *k*_0,sev_. (E) Heterogeneity of the network quantified as a coefficient of variation in actin density in the y direction. (F) Retrograde flow speed for different *k*_0,sev_. With intermediate values of *k*_0,sev_, network was more homogeneous, and the flow was faster with smaller fluctuations.

A variation in *λ*_sev_ led to similar results (Fig. S6). With higher *λ*_sev_, the severing rate becomes more sensitive to a change in local bending angles on F-actins induced by buckling.

When *λ*_sev_ was very high, severing occurred even by a small increase in the bending angles, and excessively frequent severing led to results similar to those with high *k*_0,sev_ (Figs. S6B, D-G). When *λ*_sev_ was small, F-actin severing hardly took place, resulting in the stalling of network contraction and the loss of connection between the network and the motors similar to cases with *k*_0,sev_ = 0 (Figs. S6A, C, E-G). In sum, network disassembly mediated by steady actin depolymerization and angle-dependent severing should take place at moderate rates for maintaining the dynamic steady state.

### A minimal number of motors are required for a continuous actin retrograde flow against friction from FAs

In this model, network contraction induced by motors located near the -y boundary is a main driving factor for the retrograde flow. We probed the impact of a variation in motor density (*R*_M_). An insufficient number of motors (*R*_M_ ≤ 0.003) resulted in a negligible retrograde flow and minimal network contraction because forces generated by the motors were not large enough to overcome friction exerted by FAs (Figs. 5F, G and S5C). Since network disassembly continuously occurred near the -y boundary, the stationary network eventually lost connection to motors (Figs. 5A, C, E). When *R*_M_ was increased from 0.003 to 0.005, the flow speed was abruptly increased with noticeable network contraction, implying that motor-generated forces became larger than the frictional forces exerted by FAs (Figs. 5F, G and S5C). A further increase in the number of motors (*R*_M_ ≥ 0.005) still led to a steady flow, but F-actins began to accumulate towards the -y boundary because the network contraction was faster than network disassembly occurring near the -y boundary, which is reminiscent of the formation of actin arcs (Figs. 5B, D, E, F). We measured a total force transmitted from the network to the underlying substrate by summing forces acting on individual FAs. The total force was proportional to *R*_M_ up to *R*_M_ = 0.005 and became independent of *R*_M_ at *R*_M_ > 0.005 because maximum forces that FAs can transmit are limited by the force-dependent unbinding of links forming FAs (Fig. 5G). When *R*_M_ was increased beyond 0.005, the frictional forces were relatively constant, but forces generated by motors increased, resulting in higher flow speed.

**Figure 5.**
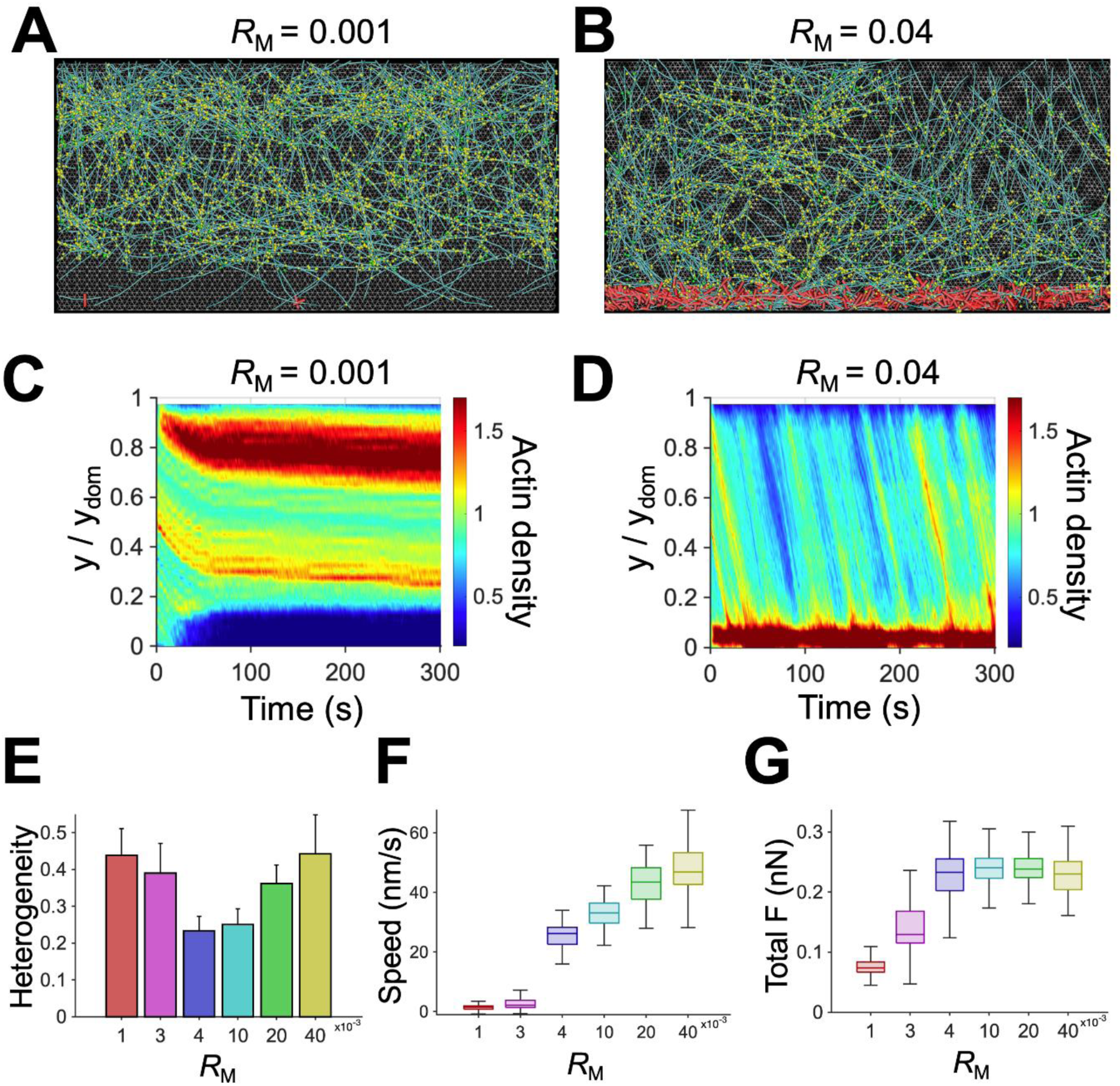
An increase in motor density makes the retrograde flow faster by enhancing contractile forces. (A, B) Snapshots of the branched network taken at ∼150 s with a lower or higher value of motor density (*R*_M_) relative to that of the reference condition, 0.004. (C, D) Kymographs of actin density as a function of y position and time with different values of *R*_M_. The case with insufficient motors did not show a noticeable flow because it could not overcome frictional forces from FAs. As *R*_M_ increased, more F-actins were accumulated as a bundle near the -y boundary. (E) Heterogeneity of the network quantified as a coefficient of variation in actin density in the y direction. The network was most homogeneous with intermediate values of *R*_M_. (F) Actin retrograde flow speed depending on *R*_M_. With higher *R*_M_, the flow tended to be faster. (G) Total force acting on the entire substrate via elastic links with different *R*_M_. The total force showed a clear plateau at high *R*_M_.

In addition to motor density, we probed the effect of the ATP-dependent unbinding rate (*k*_20_) of myosin heads which is one of the mechanochemical rates used in the parallel cluster model (48, 49). The walking and unbinding rates of motors in our model tend to be proportional to *k*_20_. With low *k*_20_, the network failed to reach a steady state because the motors walk along F-actins more slowly, resulting in lower flow speed (Figs. S7A, C, E-H). Beyond threshold walking speed (*k*_20_ ≥ 6), the network reached steady state, and the network heterogeneity noticeably decreased (Figs. S7B, D-H). With *k*_20_ = 17 s^-1^ used for physiologically relevant walking speed of non-muscle myosin II (120 nm/s) (50, 51) in the reference condition, the network showed stable flow speed and relatively homogenous morphology (Figs. S7E-H). In sum, a sufficient number of motors are necessary to overcome resistances from FAs and induce a steady retrograde flow, and with more, faster motors, the flow becomes faster, leading to F-actin accumulation into a bundle.

### Large frictional forces from FAs can frustrate the retrograde flow

In our model, FAs can be formed only in a specific region which spans across the entire domain in the x direction but occupies only a fraction of the domain in the y direction (Fig. 1C). We varied the size of the FA region by altering its fraction relative to the entire domain size (*A*_FA_) to investigate the effects of frictional forces from FAs on the dynamic steady state. Without FA region (*A*_FA_ = 0), the retrograde flow speed was the highest, and the network was slightly accumulated near the -y boundary (Figs. 6A, C, E-G and S5D), similar to observations in the case with high *R*_M_. Consistent with experimental results (52, 53), larger FA region resulted in a slower retrograde flow (Figs. 6B, D-G and S5D). With the largest FA region that we tested (*A*_FA_ = 0.5), the network was stalled at the FA region and eventually disconnected from motors, resulting in higher heterogeneity and lower flow speed. This is similar to observations made in the case with very low *R*_M_. A total force transmitted between the network and the underlying substrate increased in proportion to *A*_FA_ (Fig. 6G), with a slight decrease for the case with *A*_FA_ = 0.5 which failed to reach a steady state. When the total force was close to maximum forces generated by motors (*A*_FA_ > 0.4), the retrograde flow was frustrated and disappeared after ∼100 s (Figs. 6F, G and S5D). Overall, an increase in *A*_FA_ led to results similar to those resulting from a decrease in *R*_M_ because the retrograde flow originates from a competition between force generation from motors and frictional resistances from FAs. If contractile forces are greater than frictional forces, a steady flow emerges, and network connectivity is maintained well. If not, the flow does not appear, and the network can undergo structural discontinuity.

**Figure 6.**
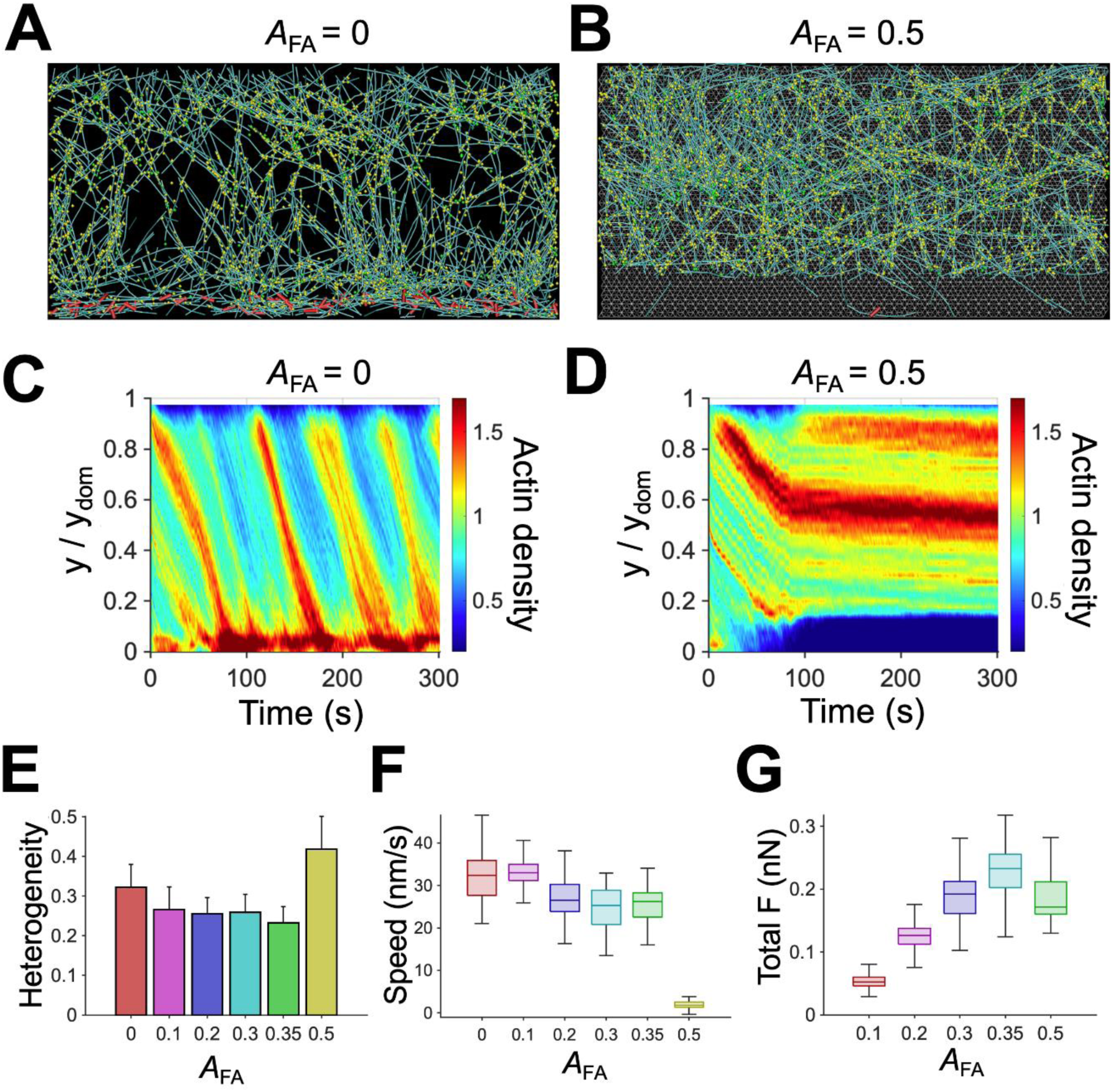
Focal adhesions (FAs) hinder the actin retrograde flow by exerting frictional forces. (A, B) Snapshots of the branched network taken at ∼150 s with smaller or larger size of the FA region (*A*_FA_) relative to that of the reference condition, 0.35. (C, D) Kymographs of actin concentration depending on time and y position with *A*_FA_. Without the FA region, the accumulation of F-actins into a bundle was noticeable, whereas large FA region led to discontinuity between the network and the motors. (E) Heterogeneity of the network quantified as a coefficient of variation in actin density in the y direction. (F) Actin retrograde flow speed depending on *A*_FA_. The flow speed tended to be smaller with larger FA region due to higher frictional forces. (G) Total force acting on the substrate by the network with different *A*_FA_. At *A*_FA_ ≤ 0.35, the total substrate force was proportional to *A*_FA_ due to the formation of more links, but the case with *A*_FA_ = 0.5 showed a lower total force because of the discontinuity between the network and the motors occurring at later times.

### Various dynamic steady states can exist

We have presented results from simulations with a variation in only one parameter relative to the reference case to demonstrate under what condition the dynamic steady state is maintained or disrupted. Our results suggest that the balance between network assembly and disassembly and the competition between contractile forces and frictional forces lead to the dynamic steady state characterized by the continuous retrograde flow. It is likely that there can be various dynamic steady states with different retrograde flow speed as long as minimum conditions for the dynamic steady state can be satisfied. Indeed, it has been experimentally observed that the lamellipodia exhibited different states and switched between them to adapt to varying environments. For example, a branched actin network underwent significant structural changes in response to mechanical loads transmitted from the extracellular environment (54, 55). We explored the possibility of different modes of dynamics steady states. Based on the observation that a decrease in *R*_M_ has opposite effects on flow speed and network contraction to those of a decrease in *A*_FA_ (Figs. 5 and 6), we first investigated whether a decrease in *A*_FA_ can rescue the case that failed to reach a steady state due to very low *R*_M_ = 0.0015. When the FA region was reduced from *A*_FA_ = 0.175 to *A*_FA_ = 0, a steady and fast retrograde flow emerged because contractile forces generated by motors could be used solely for network contraction without a competition against frictional forces exerted by FAs (Figs. 7A, C). Alternatively, the case that failed to show the actin retrograde flow due to high *A*_FA_ could be rescued by increasing *R*_M_ from 0.0015 to 0.004 (Figs. 7A, C).

**Figure 7.**
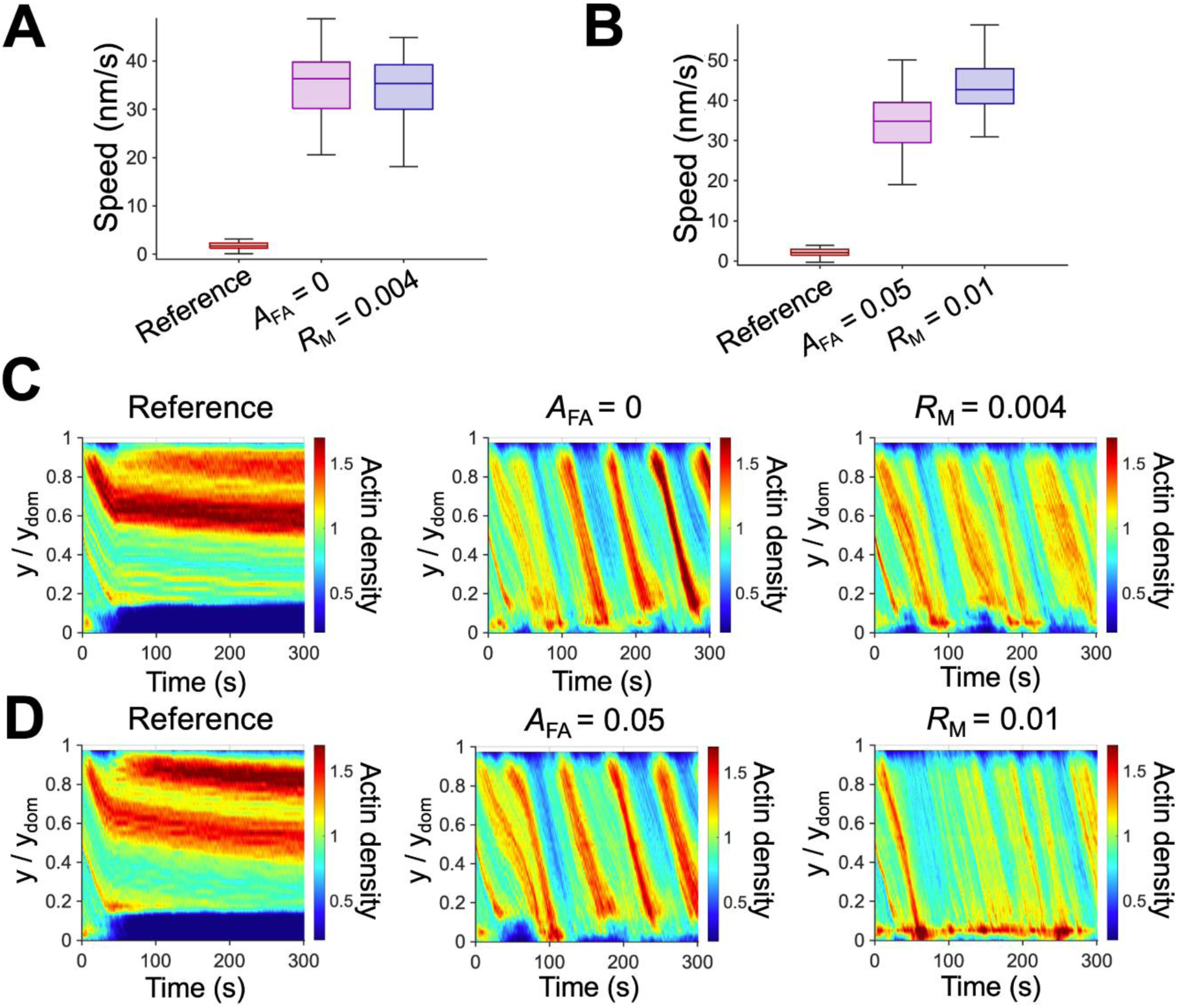
Different modes of dynamic steady states exist. (A) Two ways to rescue the case showing a negligible flow with *R*_M_ = 0.0015, *A*_FA_ = 0.175, and *k*_-,A_ = 6 s^-1^. Since contractile forces were insufficient to overcome the frictional forces in this case, it was able to be rescued by either increasing *R*_M_ to 0.004 or reducing *A*_FA_ to 0. (B) Rescuing a case exhibiting a minimal flow with *R*_M_ = 0.004, *A*_FA_ = 0.15, and *k*_-,A_ = 10 s^-1^. Since actin depolymerization was too fast compared to flow speed, this case was able to be rescued by enhancing flow speed, either by increasing *R*_M_ to 0.01 or decreasing *A*_FA_ to 0.05. (C, D) Kymographs of actin density as a function of y position and time for cases shown in (A) and (B), respectively.

We previously showed that fast depolymerization (*k*_-,A_ = 10 s^-1^) resulted in the loss of network connectivity near the -y boundary due to excessively fast network disassembly, preventing motors from binding and properly walking along F-actins. We next tested whether increasing the flow speed to match the fast depolymerization rate can rescue the case. We found that a steady flow was restored with either lower *A*_FA_ or higher *R*_M_ (Figs. 7B, D), showing how the competition between contractile and frictional forces is related to the balance between network assembly and disassembly. A notable difference is that the restored flow speed was considerably faster than that observed in the reference case. A decrease in the frictional forces or an increase in contractile forces caused the network to flow more readily towards the -y boundary, which supplied enough F-actins for motors to bind and walk along before former F-actins were completely disassembled. All of these results suggest that different types of dynamic steady states exist and a switch between these states is possible by adjusting the appropriate parameters.

## DISCUSSION

Lamellipodia, sheet-like cell protrusions, have been extensively studied during recent decades. Although the underlying molecular players and working principles for lamellipodia have been identified, it remains unclear how the dynamic steady state is maintained despite environmental changes. In this study, we simulated the branched actin network in the lamellipodia to define the mechanism of adaptive maintenance of the actin retrograde flow. We succeeded in reproducing a steady-state retrograde flow driven by actin dynamics and motor activity against FAs (Fig. 2). Through parametric studies, we showed the importance of a balance between network assembly near the leading edge and network disassembly at the rear end of the lamellipodia.

First, we found that sufficiently fast actin polymerization and high Arp2/3 density enabled the network to maintain a continuous retrograde flow by timely creating new branches near the leading edge (Figs. S2 and S3). When branch formation was delayed due to either slow actin polymerization or insufficient Arp2/3 complexes, network assembly was unable to keep up with network disassembly, resulting in network discontinuity. We also showed that a large number of ACPs could lead to the uniform contraction of an entire network toward the rear as a single entity by enhancing network connectivity (Fig. S4). This is consistent with our previous study which showed that many cross-linking points between F-actins were required for global network contraction into a single cluster (56). In the lamellipodia, new branches are formed toward the leading edge with the characteristic angle of 70°, so it is hard to expect that the network has percolation in a direction perpendicular to the flow direction. Thus, the presence of ACPs is crucial for contracting the network as a whole. A recent filament-level computational model for branched actin networks also included ACPs for achieving sufficient network connectivity (45).

Network disassembly at the rear in our model takes place in two different ways: F-actin depolymerization and severing. We found that these two dynamic events should take place at moderate rates at the rear to induce a steady retrograde flow without structural discontinuity or F-actin accumulation (Figs. 3 and 4). In our model, we allowed F-actin depolymerization to occur only at the rear. It was experimentally shown that the depolymerization rate gradually increases towards the rear (57, 58), which might be attributed to ATP hydrolysis of F-actins. It has been known that ADP-F-actin is more susceptible to depolymerization and severing (59–61). We did not include a gradually changing depolymerization rate or ATP hydrolysis of actin for simplicity. Unlike depolymerization, F-actin severing can occur anywhere in our model. However, we assumed that the severing rate exponentially increases as a local bending angle on F-actin increases as in our previous studies (56, 62). This led to the dominant occurrence of F-actin severing at the rear because F-actins undergo buckling during network compaction into a bundle-like structure as we showed before (63). In our previous study, we demonstrated that F-actin severing was indispensable for reproducing the pulsatile (i.e., reversible) contraction of actomyosin cortex observed in cells because the severing can locally increase the network disassembly rate within aggregating structures by creating more pointed ends (Figs. 4 and S6) (64). Likewise, the buckling-induced severing helped the disassembly of the dense actin structure at the rear resulting from network accumulation.

Consistent with experimental studies, we found that larger force generation from more motors and less frictional forces from smaller FA region could result in a faster retrograde flow speed (Figs. 5, 6, and S7) (22). The importance of myosin motor activity and FAs has been demonstrated in many studies (65, 66). Contractile forces generated by motors contract the actin network toward the rear to drive a retrograde flow, whereas FAs resist the flow by exerting frictional forces via transient links formed between the underlying substrate and the network. Thus, the competition between the contractile forces and the frictional forces determines the nature and speed of the retrograde flow. We observed that the flow was negligible when motors could not overcome resistances from FAs. Once the contractile forces overcame the fractional forces from FAs, the flow speed abruptly increased due to frictional slippage between the network and the underlying substrate, which was shown in a previous study (66).

A recent study has shown that the mechanosensing ability of migrating cells allows them to adapt to different extracellular environments by adjusting the polymerization rate and density of F-actins in lamellipodia (67). In addition, compressive loads were found to increase the density of growing filament ends in the branched actin network at the leading edge by decreasing the capping rate, instead of directly increasing an overall nucleation rate (68). These experimental findings suggest that cells constantly adapt to time-varying environments by altering the activities and concentrations of key regulators. We showed that the branched network in lamellipodia can attain different modes of dynamic steady states (Fig. 7). Specifically, we demonstrated that cases exhibiting a negligible retrograde flow due to one parameter can be rescued by varying other parameter value. For example, the case lacking contractile forces from motors could be rescued by decreasing the FA size. The case with a negligible retrograde flow due to rapid actin depolymerization could be rescued by enhancing the flow speed by including more motors or by decreasing the FA region.

Previous computational models of the lamellipodia have provided valuable insights into understanding the role of different molecular players. The molecular clutch model was developed to explain the actin retrograde flow driven by myosin activity and actin polymerization (66). Although this one-dimensional model could capture the essential aspects of the retrograde flow, it was designed for representing filopodia, not lamellipodia. Thus, network connectivity mediated by Arp2/3 and ACPs, and network disassembly induced by buckling-induced severing could not be considered. Razbin et al. investigated the properties of a branched network in the lamellipodia, using a simple model where mother filaments grafted at one end undergo polymerization against a membrane at the leading edge (42). Hu et al. developed a model incorporating physico-chemical reaction and diffusion processes in a stochastic manner (41). Their model showed the importance of the interplay between F-actin, Arp2/3, and capping proteins for lamellipodial protrusion dynamics. However, these models aimed to probe the role of specific molecular players in protrusion dynamics rather than focusing on interactions between several key molecular players. Since they did not consider a competition between FAs and motor proteins, they could not capture the essential aspect of the lamellipodia. Schreiber et al. previously showed a three-dimensional stochastic model that reproduced the force-velocity relationship of cells observed experimentally (40). Although this model was able to account for the retrograde flow in the lamellipodia, it comprised simplified actin dynamics, and the interplay between different factors, such as ACP binding, severing, and motor activity, was not considered.

Although our model is more sophisticated than any other model developed for lamellipodia, the model still has some limitations. Our model does not consider the deformation of a cell membrane at the leading edge induced by the polymerization of branches via the Brownian ratchet model (69) or thermal fluctuation. Additionally, our model considers only myosin II as molecular motors driving the retrograde flow. Prior studies have shown that the retrograde flow persisted even after the inhibition of myosin II via blebbistatin, implying the existence of other molecular motors driving the retrograde flow, such as myosin Ic attached to the membrane, as seen in growth cones (70–72). In addition, we allowed the polymerization and depolymerization of F-actins to occur only in specific regions. In our model, the lamellipodia are considered as a closed system, whereas in vivo, the actin pool is shared with the rest of the cell. It is also important to note that to make actin dynamic more realistic, considering the explicit states of ATP-actin and ADP-actin with cofilin activity would be necessary. Lastly, the maturation of FAs is not incorporated in our model by assuming that only nascent FAs are formed between the branched network and the underlying substrate (Fig. 2B).

## CONCLUSIONS

Our computational model successfully reproduced the actin retrograde flow of the branched network observed in the lamellipodia and provided insights into understanding the characteristics of the dynamic steady state of the flow. Our results suggested the importance of i) the balance between network assembly at the leading edge and network disassembly at rear and ii) a competition between myosin-generated contractile forces and FA-induced frictional forces acting between the network and a surrounding environment. We further demonstrated that different modes of dynamic steady states are possible, which sheds light on prior experimental findings about the adaptability of the lamellipodia to different mechanical environments. In the near future, we will incorporate a simplified cell membrane that can be deformed by elongating branches to simulate cell protrusion and the retrograde flow at the same time.

## AUTHOR CONTRIBUTIONS

J.H.K. and T.K. designed the project. J.H.K. performed simulations and analyzed data obtained from the simulations. T.K. supervised all computational studies. All the authors participated in writing the manuscript.

## DATA AVAILABILITY STATEMENT

The data that support the findings of this study are available from the corresponding author upon reasonable request.

## CONFLICT OF INTEREST DISCOSURE

The authors declare no conflict of interest.

