## Supporting Information for "The Mechanism of Dynamic Steady States in Lamellipodia"

*Brownian dynamics via the Langevin equation*

In our agent-based model, cytoskeletal elements consist of segments which are defined by their endpoints. The displacements of the endpoints of all segments are updated by the Langevin equation at each time step with inertia neglected:

$$\mathbf{F}_i - \zeta_i \frac{d\mathbf{r}_i}{dt} + \mathbf{F}_i^{\mathrm{T}} = 0 \tag{S1}$$

where $\mathbf{F}_i$, $\zeta_i$, and $\mathbf{r}_i$ represent the deterministic forces, drag coefficient, and position vector of the *i*th endpoint, respectively, and $t$ is time. $\zeta_i$ is calculated via an approximated form for a cylindrical object (73):

$$\zeta_i = 3\pi\mu r_{c,i} \frac{3 + 2r_{0,i} / r_{c,i}}{5} \tag{S2}$$

where $\mu$ is the viscosity of the surrounding medium, and $r_{0,i}$ and $r_{c,i}$ are the length and diameter of segments, respectively. $\mathbf{F}^{\mathrm{T}}_i$ is a stochastic force satisfying the fluctuation-dissipation theorem (74):

$$\left\langle \mathbf{F}_i^{\mathrm{T}}(t)\mathbf{F}_j^{\mathrm{T}}(t) \right\rangle = \frac{2k_B T \zeta_i \delta_{ij}}{\Delta t} \boldsymbol{\delta} \tag{S3}$$

where $\boldsymbol{\delta}$ is a second-order tensor, $\delta_{ij}$ is the Kronecker delta, $k_B T$ is thermal energy, and $\Delta t$ = $1.15\times10^{-5}$ s is time step. The positions of the cylindrical segments are updated every time step using calculated velocities from the Langevin equation ($d\mathbf{r}_i/dt$) and the Euler integration scheme:

$$\mathbf{r}_i(t + \Delta t) = \mathbf{r}_i(t) + \frac{d\mathbf{r}_i}{dt}\Delta t = \mathbf{r}_i(t) + \frac{1}{\zeta_i}\left(\mathbf{F}_i + \mathbf{F}_i^{\mathrm{T}}\right)\Delta t \tag{S4}$$

For the deterministic forces, we consider i) extensional forces which maintain the equilibrium lengths of segments, ii) bending forces that maintain equilibrium angles formed by segments, iii) torsion forces which maintain torsional angles near their equilibrium values, and iv) repulsive forces which account for volume-exclusion effects between neighboring actin segments. The extensional, bending, and torsional forces are defined by the following potentials:

$$U_s = \frac{1}{2}\kappa_s (r - r_0)^2 \tag{S5}$$

$$U_{\mathrm{b}} = \frac{1}{2}\kappa_{\mathrm{b}}\left(\theta - \theta_0\right)^2 \tag{S6}$$

$$U_{\mathrm{t}} = \frac{1}{2}\kappa_{\mathrm{t}}\left(\phi - \phi_0\right)^2 \tag{S7}$$

where $\kappa_s$, $\kappa_b$, and $\kappa_t$ represent the extensional, bending, and torsional stiffnesses, respectively. $r$ and $r_0$ are the instantaneous and equilibrium lengths of segments, $\theta$ and $\theta_0$ are instantaneous and equilibrium angles formed by interconnected segments, and $\phi$ and $\phi_0$ are instantaneous and equilibrium torsional angles. The repulsive forces acting between F-actins are represented by the following harmonic potential (75):

$$U_{\mathrm{r,A}} = \begin{cases} \frac{1}{2}\kappa_{\mathrm{r,A}}\left(r_{12} - r_{\mathrm{c,A}}\right)^2 & \text{if} \quad r_{12} < r_{\mathrm{c,A}} \\ 0 & \text{if} \quad r_{12} \geq r_{\mathrm{c,A}} \end{cases} \tag{S8}$$

where $\kappa_{r,A}$ represents the strength of repulsive forces, $r_{12}$ is a distance between two neighboring elements, and $r_{c,A}$ is the diameter of actin segments.

*Mechanics of cytoskeletal components*

The extensional ($\kappa_{s,A}$) and bending ($\kappa_{b,A}$) stiffnesses of actin segments maintain their equilibrium length ($r_{0,A}$ = 140 nm) and equilibrium angle ($\theta_{0,A}$ = 0 rad), respectively (Fig. S1). The equilibrium length of each ACP segment ($r_{0,ACP}$ = 20 nm) and an equilibrium angle between two ACP segments ($\theta_{0,ACP}$ = 0 rad) are maintained by extensional ($\kappa_{s,ACP}$) and bending ($\kappa_{b,ACP}$) stiffnesses, respectively.

Arp2/3 is involved with one extensional stiffness, four bending stiffnesses, and one torsional stiffness. The equilibrium length of each Arp2/3 segment ($r_{0,Arp2/3}$ = 35 nm) is maintained by extensional stiffness ($\kappa_{s,Arp2/3}$). An equilibrium angle between two Arp2/3 segments ($\theta_{0,Arp2/3}$ = 0 rad) is maintained by bending stiffness ($\kappa_{b,Arp2/3c}$). Two bending stiffnesses ($\kappa_{b,Arp2/3m}$ and $\kappa_{b,Arp2/3d}$) maintain an equilibrium angle between one segment of Arp2/3 and the mother filament ($\theta_{0,Arp2/3m}$ = 90° = 1.57 rad) and an equilibrium angle between the other segment of Arp2/3 and the daughter filament ($\theta_{0,Arp2/3d}$ = 20° = 0.35 rad), respectively. Another bending stiffness ($\kappa_{b,Arp2/3f}$) maintains an equilibrium angle between the mother and daughter filaments ($\theta_{0,Arp2/3d}$ = 70° = 1.22 rad). Torsional stiffness ($\kappa_{t,Arp2/3}$) maintains a zero torsional angle between the mother and daughter filaments to enforce them to exist on a single plane.

The equilibrium length of motor backbone segments ($r_{s,MB}$ = 42 nm) and an equilibrium angle between the backbone segments ($\theta_{s,MB}$ = 0 rad) are maintained by bending ($\kappa_{b,MB}$) and extensional ($\kappa_{s,MB}$) stiffnesses, respectively. The value of $\kappa_{s,MB}$ is equal to $\kappa_{s,A}$, whereas the value of $\kappa_{b,MB}$ is much greater than $\kappa_{b,A}$. The extension of each motor arm is regulated by the two-spring model with the stiffnesses of transverse ($\kappa_{s,M1}$) and longitudinal ($\kappa_{s,M2}$) springs. The transverse spring maintains an equilibrium distance ($r_{0,M1}$ = 10 nm) between the endpoint of a motor backbone and an actin segment where the arm of the motor binds, and the longitudinal spring helps maintain the right angle between the motor arm and the actin segment ($r_{0,M2}$ = 0 nm).

*Dynamic behaviors of cytoskeletal components*

F-actin can undergo de novo nucleation, polymerization, depolymerization, and angle-dependent severing. The de novo nucleation occurs via the appearance of a single actin segment with a constant rate constant, $k_{n,A}$. Actin polymerization occurs only from the barbed end of F-actin with a constant rate constant, $k_{+,A}$, by adding one segment. Actin depolymerization takes place only from the pointed end of F-actin at a constant rate, $k_{-,A}$, by removing one segment. Both polymerization and depolymerization occur without dependence on force. F-actin is not allowed to elongate beyond 0.98 μm to mimic the activity of capping proteins (76, 77). F-actin severing occurs with a rate, $k_{sev}$, proportional to a local bending angle by removing one segment at the mid of F-actin:

$$k_{sev} = k_{0,sev} \exp\left(\lambda_{sev}\theta\right) \quad \text{(S9)}$$

where $k_{0,sev}$ and $\lambda_{sev}$ represent the zero-angle severing rate constant and angle sensitivity, respectively (78).

ACPs connect pairs of F-actins to form functional cross-linking points. They bind to binding sites located on F-actins every 7 nm first with a constant rate constant, $k_{+,ACP}$, and then bind to binding sites on the other F-actin with the same rate constant if they are sufficiently proximal to each other. Unlike our previous studies (63, 79), it is assumed that ACPs cannot unbind from F-actins by themselves, meaning that they form permanent cross-links.

Arp2/3 binds to binding sites on F-actins with a constant rate constant, $k_{+,Arp2/3}$. Immediately after binding, it nucleates a daughter filament; a new actin segment appears with the characteristic angle of 70° relative to the mother filament with a directional bias towards the leading edge. Connections between Arp2/3 and mother/daughter filaments are assumed to be permanent.

Motor arms bind to binding sites on actin segments with the rate of $40N_h$ where $N_h$ is the number of myosin heads represented by one motor arm. After binding, motor arms are able to walk along F-actin at a force-dependent rate, $k_{w,M}$, and unbind from actin segments at a force-dependent rate, $k_{u,M}$. $k_{w,M}$ and $k_{u,M}$ are determined by the parallel cluster model to account for the mechanochemical cycle of non-muscle myosin II (11, 12, 48, 49). Details of implementation and benchmarking of the parallel cluster model were described in our previous study (80). The parallel cluster model assumes three states of myosin heads and uses 5 transition rates between those states. One of them is the ATP-dependent unbinding rate of myosin heads ($k_{20}$) which defines a transition rate from the post power stroke state to the unbound state. $k_{w,M}$ and $k_{u,M}$ in the model are lower with smaller $k_{20}$ or with a larger applied load due to an assumption that motors exhibit a catch-bond behavior. The unloaded walking velocity and stall force of motor arms with the reference value of $k_{20}$ are set to 120 nm/s and 5.33 pN, respectively. If all arms of a motor lose connection to F-actin, its backbone is disassembled instantaneously and assembled on F-actin in a different location. The backbone assembly cannot occur if there is no F-actin to bind within the motor region. Thus, the number of motors may look different on snapshots between cases even if the motor density is identical.

In case of depolymerization or severing of F-actin, Arp2/3, ACPs, and motor arms bound to a disappearing actin segment lose one connection to F-actin. Then, they are able to bind to a new binding site on F-actin.

*Models for a substrate and nascent focal adhesions*

A substrate underlying the network is coarse-grained into a two-dimensional triangulated mesh (Fig. 1B). Extensional stiffness ($\kappa_{s,sub}$) maintains the length of chains between mesh nodes near their equilibrium length ($r_{0,sub}$ = 50 nm). Bending stiffness ($\kappa_{b,sub}$) maintains an angle formed by mesh nodes near its equilibrium value ($\theta_{0,sub}$ = 60° = 1.05 rad). The movement of mesh nodes in the $z$ direction is not allowed, constraining the substrate as a plane located at $z = 0$. In our previous study (18), we have explored the effect of a variation in either of $\kappa_{s,sub}$, $\kappa_{b,sub}$, or $r_{0,sub}$ on the substrate stiffness, $\kappa_{eff}$. $\kappa_{eff}$ was found to be proportional to $\kappa_{s,sub}$ and $\kappa_{b,sub}$ but inversely proportional to $r_{0,sub}$. With parameter values that we use in this study ($\kappa_{s,sub} = 1.0\times10^{-4}$ N/m, $\kappa_{b,sub} = 2.4\times10^{-20}$ Nm, $r_{0,sub} = 5.0\times10^{-8}$ m), $\kappa_{eff}$ is ~$2.4\times10^{-4}$ N/m. Thus, the substrate used in this model is considered rather soft.

To mimic the formation of nascent FAs, each mesh node is able to form a single link with any endpoint of actin segments if they are located within 200 nm. Then, this link behaves as an elastic linear spring whose equilibrium length is equal to the initial length of the link. Extensional stiffness ($\kappa_{s,C}$) maintains the equilibrium length of the link, and each link is considered as a nascent FA site, hindering the retrograde flow caused by motor activity. These links can break at a force-dependent rate, $k_{u,C}$:

$$k_{u,C} = \begin{cases} k_{u,C}^{0} \exp\left( \dfrac{\lambda_{u,C} \left| \mathbf{F}_{s,C} \right|}{k_B T} \right) & \text{if } r \geq r_{0,C} \\ k_{u,C}^{0} & \text{if } r < r_{0,C} \end{cases} \tag{S10}$$

where $k^{0}_{u,C}$ and $\lambda_{u,C}$ represent the zero-force unbinding rate constant and force sensitivity, respectively, and $\mathbf{F}_{s,C}$ is a spring force acting on the link. $r$ and $r_{0,C}$ are instantaneous and equilibrium lengths of the link, respectively.

*Computational setup*

To simulate the sheet-like, flat geometry of lamellipodia, a relatively wide and thin three-dimensional domain with the dimensions of 5×2.5×0.1 μm in x, y, and z directions is employed. Periodic boundary conditions are applied in the x direction, whereas boundaries normal to the y and z directions exert repulsive forces to elements that are displaced out of the domain. The +y boundary represents the leading edge, and the -y boundary corresponds to the interface between lamellipodia and lamella. We do not consider a movable membrane on the leading edge for simplicity. A triangular mesh is created at z = 0 as an underlying substrate. A branched network is formed via self-assembly of F-actin, Arp2/3, and ACP, with the barbed ends of F-actins biased toward +y direction. Specifically, the actin network is created from the -y boundary via the de novo nucleation of seed filaments with their barbed ends directed toward the +y direction. F-actins are elongated up to 0.98 μm via polymerization and form branches via the side binding of Arp2/3. Only ~90% of actin segments are used for network assembly to leave some free segments. ACPs physically interconnect F-actins to increase network connectivity. To reflect the primary location of myosin motors at the interface between the lamellipodia and lamella (31), we specified a region near the -y boundary (y = 0 – 0.1875 μm) for motor activity with binding, walking, and unbinding. To prevent motors from drifting toward the +y boundary, we reduced the mobility of motors by increasing their drag coefficient 1000-fold relative to a value calculated via Eq. S2.

After network formation, motors walk along F-actins toward the barbed end to contract the network toward the -y boundary. To allow for a continuous actin retrograde flow, F-actins are depolymerized from their pointed ends in a region near the -y boundary (y = 0 – 0.375 μm), whereas they are polymerized from their barbed ends in a region near the +y boundary (y = 2.125 – 2.5 μm). Free actin segments generated from depolymerization and severing of F-actins are used for polymerization at the leading edge. Reference parameter values are selected to reproduce physiologically relevant retrograde flow speed. Under the reference condition, actin concentration ($C_A$) is 250 μM, ACP density ($R_{ACP}$) is 0.04, Arp2/3 density ($R_{Arp2/3}$) is 0.01, and motor density ($R_M$) is 0.004. Note that for computational efficiency, actin concentration of 250 μM we use in this study is lower than ~500 μM measured in the real lamellipodia via electron microscopy (81). We performed simulations for 300 s with a change in parameter values. The reference values for all parameters are listed in Table S1.

*Measurement, analysis, and network visualization*

The retrograde flow speed is calculated every 2 s. The calculation is done using the y component of the velocities of F-actins located within a rectangular region near the +y boundary (Fig. 1C).

F-actin density is calculated every 1 s. First, the domain is divided into 40 subdomains in the y direction. In each subdomain, the number of actin segments is divided by $N_A$/40, where $N_A$ is the total number of actin segments in an entire domain. If actin segments are uniformly distributed, this normalized value would be close to 1. The heterogeneity of the network morphology was quantified by the coefficient of variation of F-actin density in the y direction. The standard deviation of F-actin density was calculated for each of the 40 subdomains, then divided by the mean and averaged over the simulation time; a lower value from this calculation indicates a more homogenous network.

Forces acting between the substrate and the network are calculated using links forming between membrane nodes and F-actins. Using the equilibrium length and spring constant of links (Table S1), individual forces acting on each nascent FA are calculated in each time step. The total substrate force is calculated by summing up forces acting on all links.

F-actin, Arp2/3, ACP, motor, and underlying substrate are visualized using Visual Molecular Dynamics (VMD, University of Illinois at Urbana-Champaign).

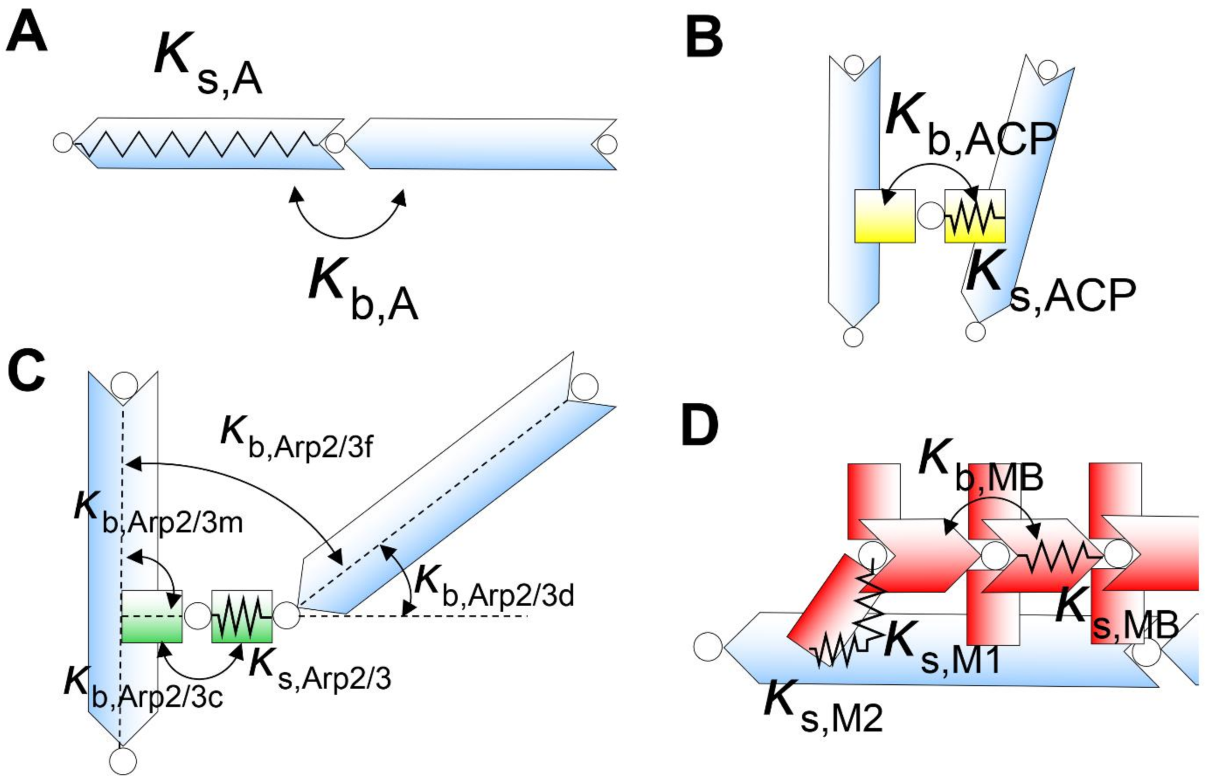


**Figure S1. Schematics showing stiffness parameters for each cytoskeletal component in the model.** (A) F-actin, (B) ACP, (C) Arp2/3 complex, and (D) motor. $\kappa_b$ and $\kappa_s$ indicate bending and extensional stiffnesses that maintain equilibrium angles and lengths, respectively. In (C), the torsional stiffness is not included in the schematic. Detailed descriptions about these stiffness parameters and their values are written in Supporting Information and Table S1.

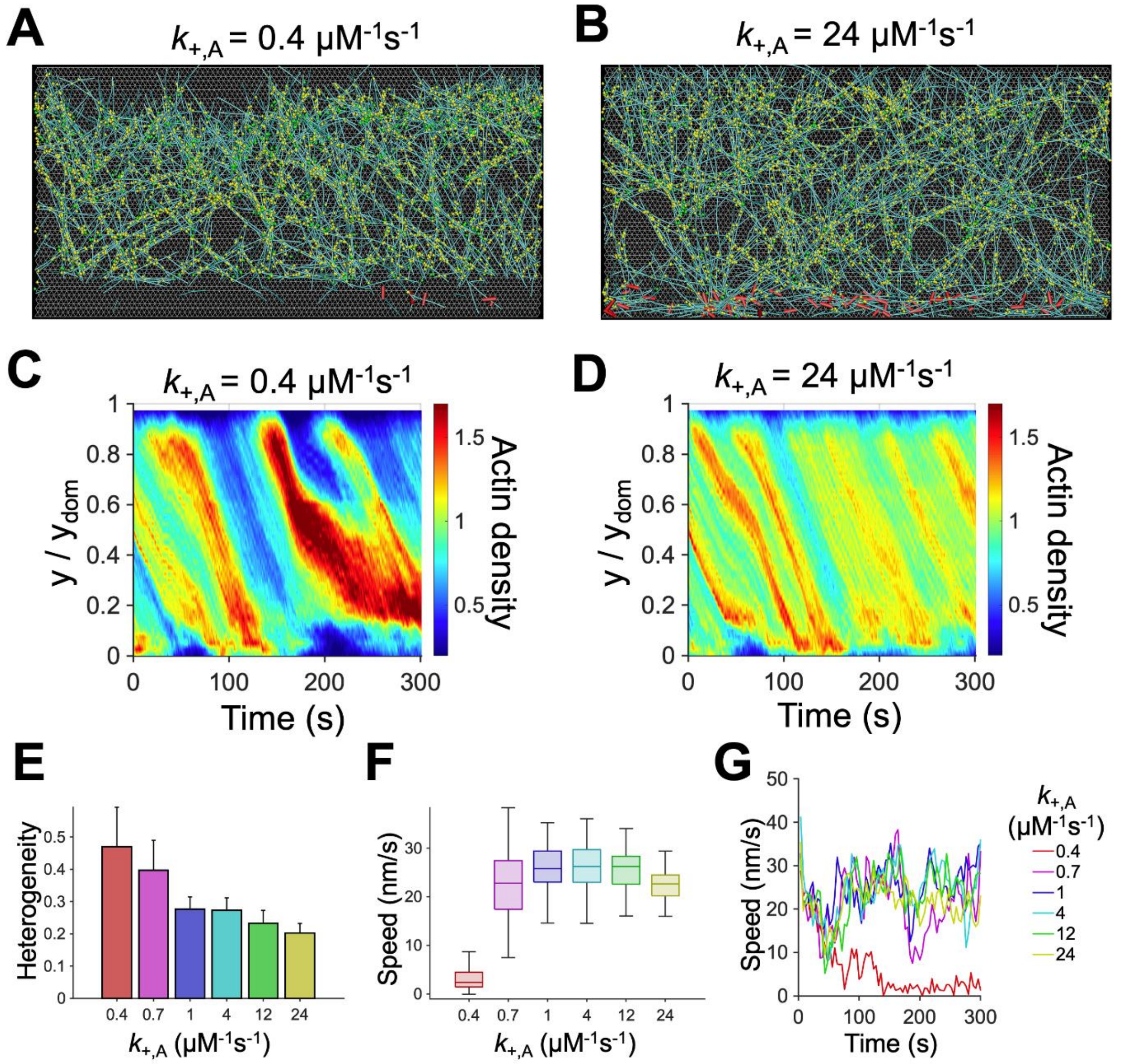


**Figure S2. Deficient actin polymerization disrupts a continuous retrograde flow.** (A, B) Snapshots of the branched network taken at ~150 s with a lower or higher polymerization rate constant ($k_{+,A}$) than that of the reference case, 12 μM$^{-1}$s$^{-1}$. (C, D) Kymographs of actin concentration as a function of y position and time with different $k_{+,A}$. With lower $k_{+,A}$, network heterogeneity increased because the network was contracted toward the -y boundary before sufficient F-actins were assembled near the +y boundary. (E) Heterogeneity of the network quantified as a coefficient of variation in actin density in the y direction. The network was more heterogeneous with deficient actin polymerization. (F) Retrograde flow speed depending on $k_{+,A}$. With deficient actin polymerization, flow speed is lower, and the network failed to reach steady state. (G) Time evolution of retrograde flow speed.

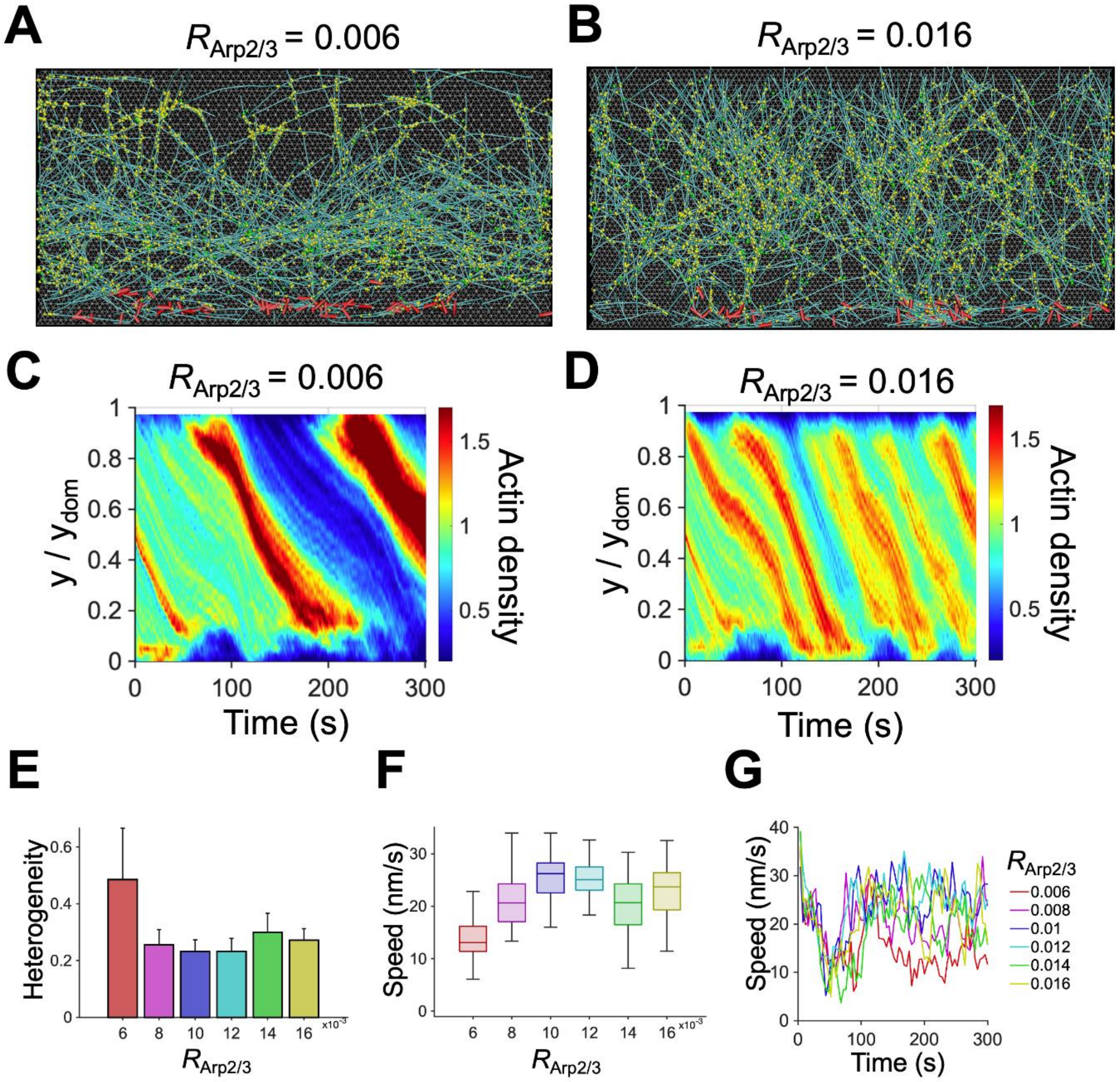


**Figure S3. Sufficiently high Arp2/3 density is required to maintain the continuity of branched network architecture.** (A, B) Snapshots of the branched network taken at ~150 s with a lower or higher Arp2/3 density ($R_{\mathrm{Arp2/3}}$) relative to that of the reference condition, 0.01. (C, D) Kymographs of actin concentration as a function of y position and time with different $R_{\mathrm{Arp2/3}}$. With low $R_{\mathrm{Arp2/3}}$, the network could not grow sufficiently, leading to higher network heterogeneity and lack of continuity in the y direction. By contrast, high $R_{\mathrm{Arp2/3}}$ resulted in the formation of more branches on a fraction of vertically growing structures, so the network became more heterogeneous with lower connectivity in the x direction. (E) Heterogeneity of the network quantified as a coefficient of variation in actin density in the y direction. (F) Retrograde flow speed with different $R_{\mathrm{Arp2/3}}$. A decrease in the flow speed was noticeable at $R_{\mathrm{Arp2/3}} < 0.01$. (G) Time evolution of retrograde flow speed.

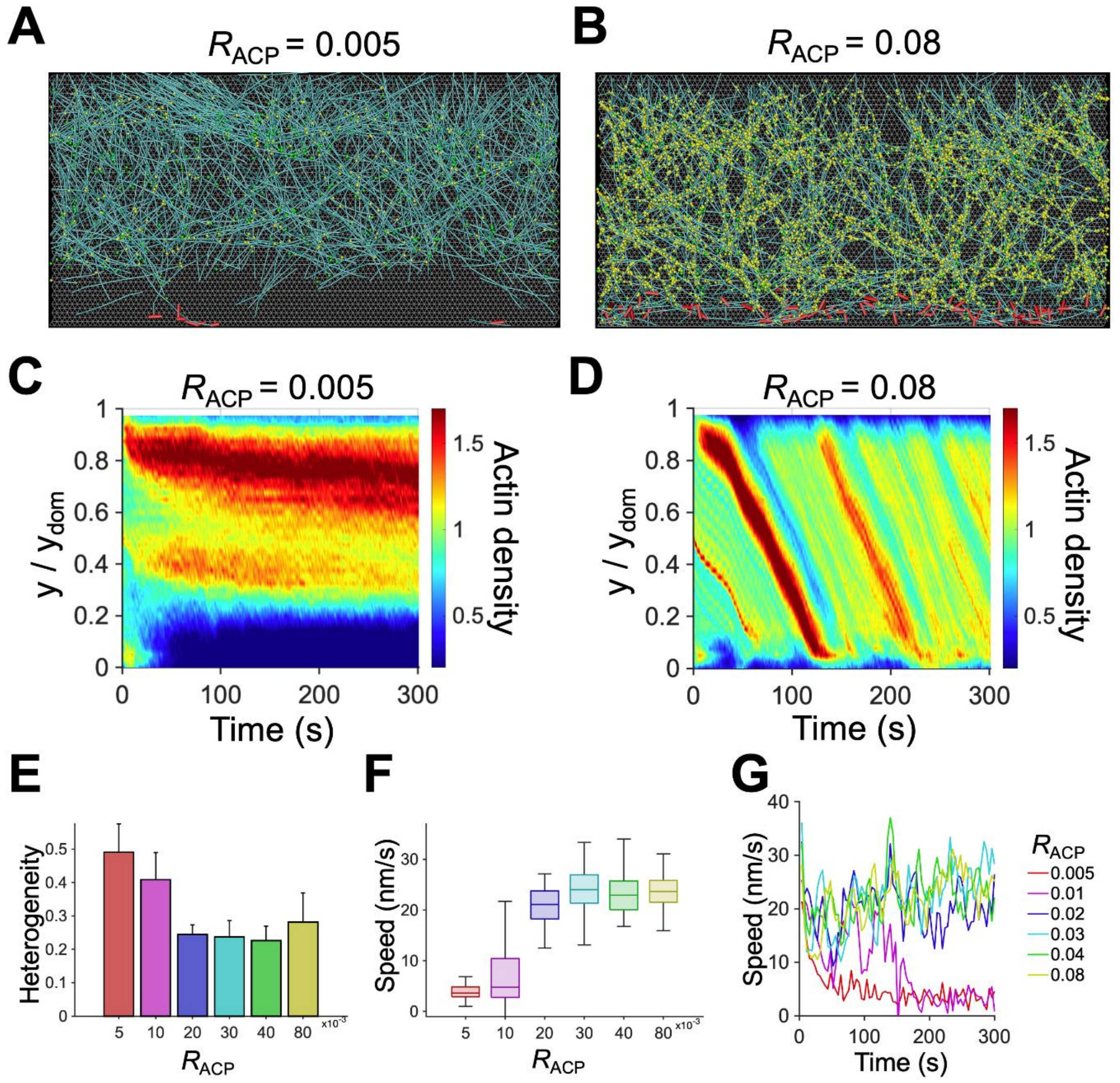


**Figure S4. More actin crosslinking proteins (ACPs) enhance network connection and thus lead to a continuous flow and homogeneous network morphology.** (A, B) Snapshots showing the branched network taken at ~150 s with a lower or higher $R_{ACP}$ relative to that of the reference condition, 0.04. (C, D) Kymographs of actin concentration as a function of y position and time with different $R_{ACP}$. With higher $R_{ACP}$, the network showed more homogeneous morphology and a continuous flow. (E) Heterogeneity of the network quantified as a coefficient of variation in actin density in the y direction. (F) Retrograde flow speed with different $R_{ACP}$. With low $R_{ACP}$ values, network heterogeneity was higher, and flow speed was slower. (G) Time evolution of retrograde flow speed.

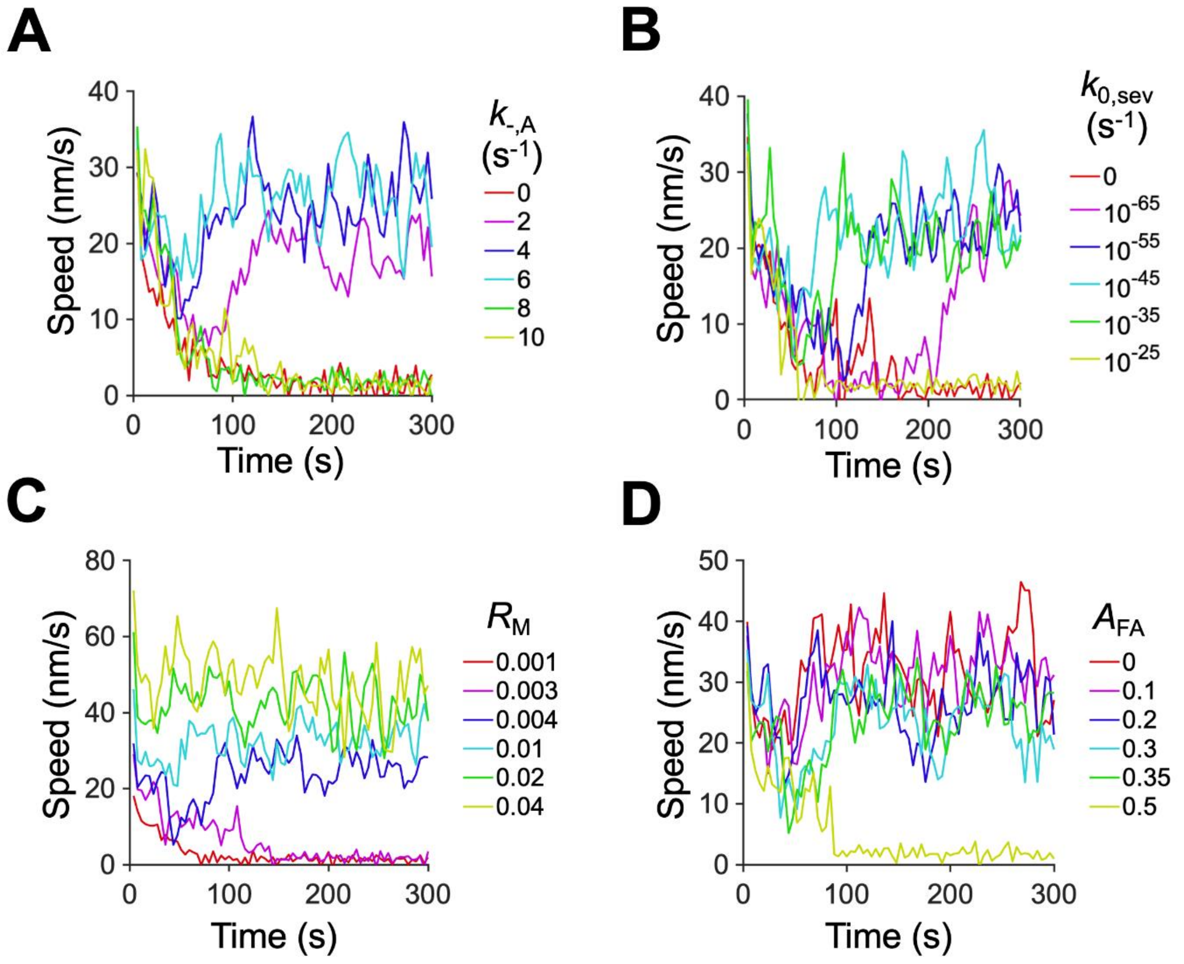


**Figure S5. Time evolution of retrograde flow speed with a change in various parameters**. (A) Actin depolymerization rate ($k_{-,A}$). (B) Actin severing rate constant ($k_{0,sev}$). (C) Motor density ($R_M$). (D) Relative size of FA region ($A_{FA}$).

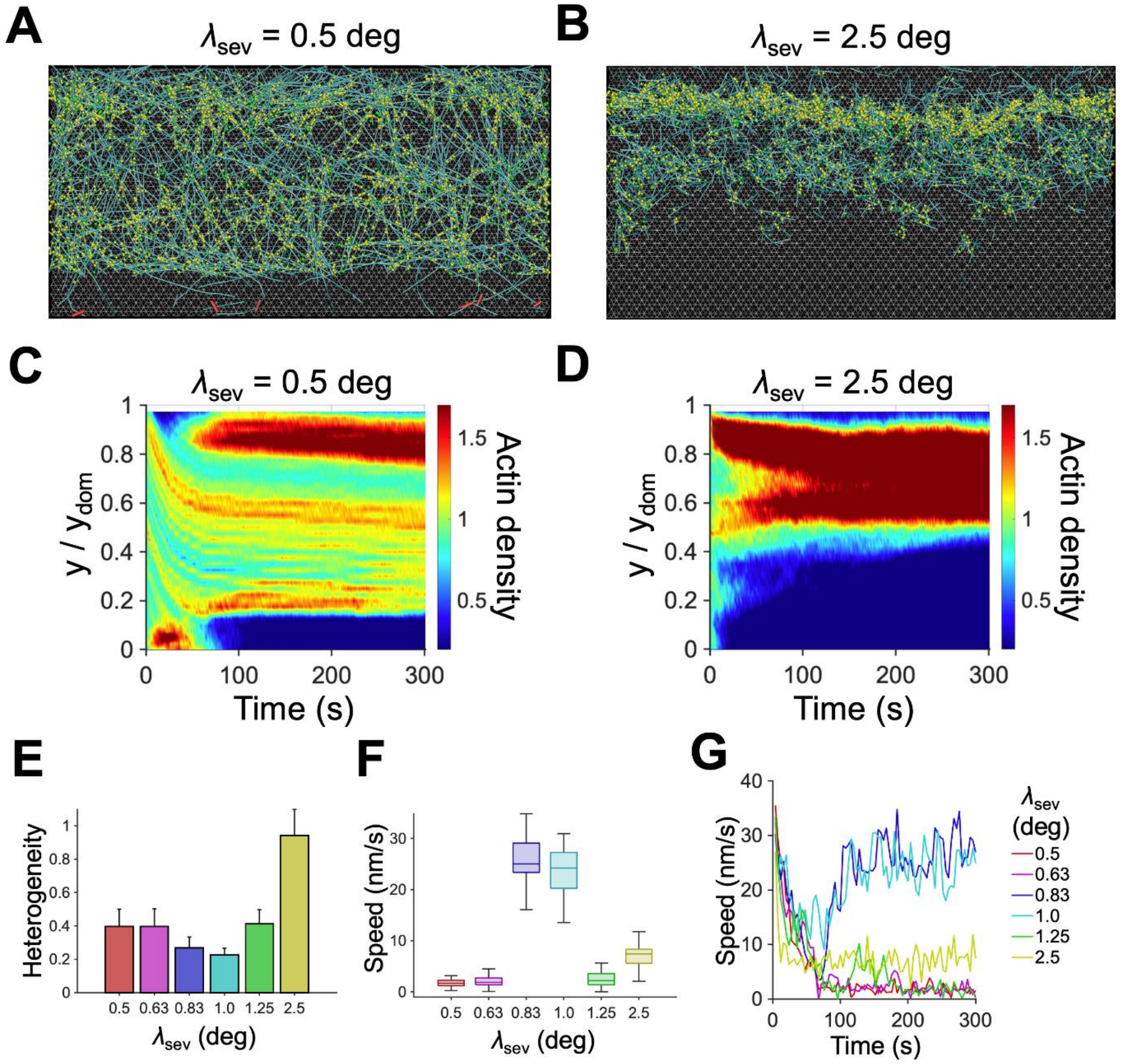


**Figure S6. Effects of the angle sensitivity of F-actin severing ($\lambda_{sev}$).** (A, B) Snapshots of the branched network taken at ~150 s with lower or higher $\lambda_{sev}$ relative to that of the reference condition, 1.0 deg. (C, D) Kymographs of actin concentration as a function of y position and time with different $\lambda_{sev}$. (E) Heterogeneity of the network quantified as a coefficient of variation in actin density in the y direction. (F) Retrograde flow speed with different $\lambda_{sev}$. Cases with intermediate $\lambda_{sev}$ reached a steady state with faster flow speed and homogenous network. (G) Time evolution of retrograde flow speed.

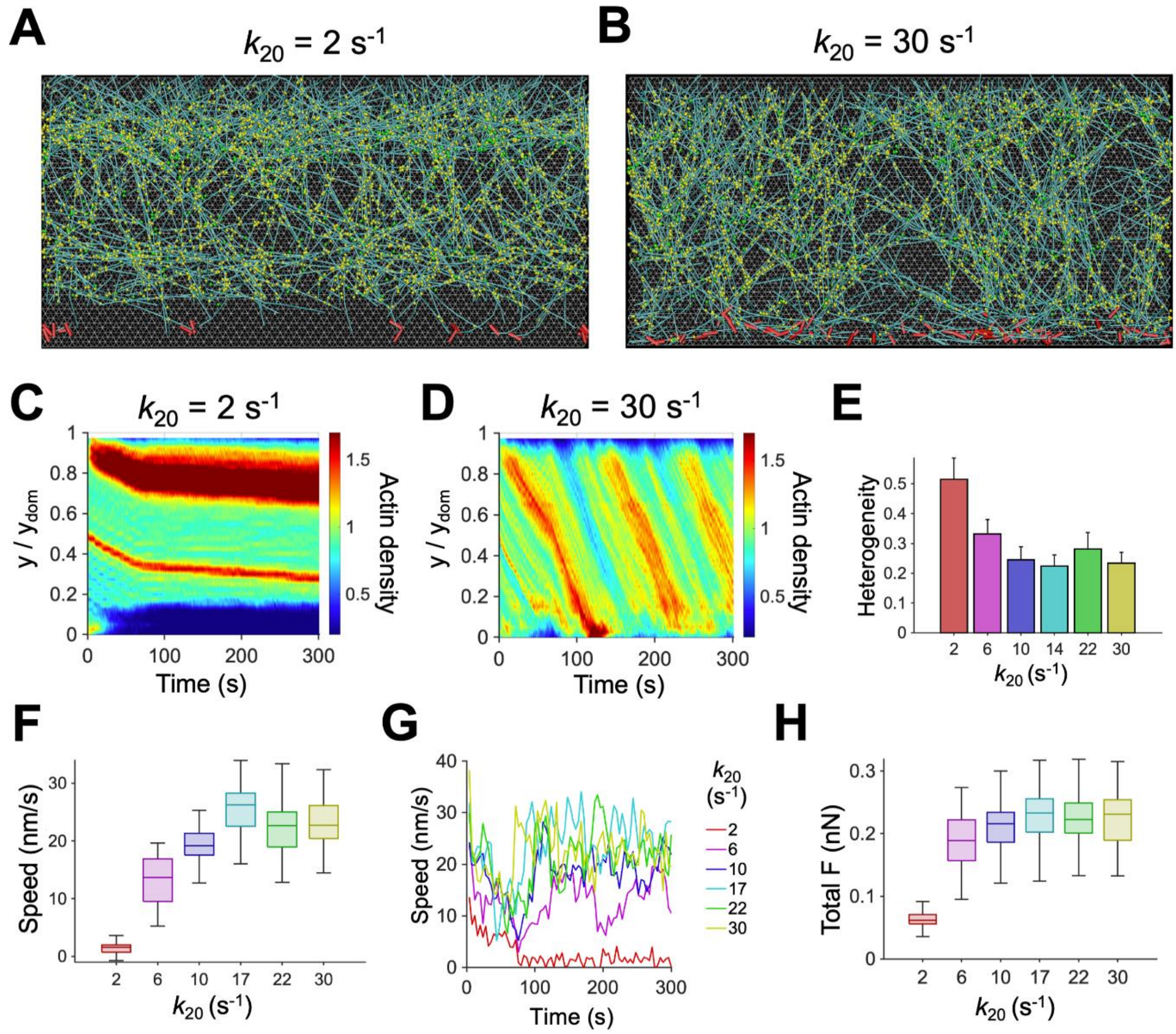


**Figure S7. Effect of the ATP-dependent unbinding rate of myosin heads ($k_{20}$) on a steady state.** With higher $k_{20}$, walking speed tends to be higher. (A, B) Snapshots of the branched network taken at ~150 s with lower or higher $k_{20}$ relative to that of the reference condition, 17 s$^{-1}$. (C, D) Kymographs of averaged actin concentration as a function of y position and time with lower or higher $k_{20}$. With the lowest $k_{20}$, the network was disconnected from motors due to excessively slow flow speed. (E) Heterogeneity of the network quantified as a coefficient of variation in actin density in the y direction. (F) Retrograde flow speed with different $k_{20}$. (G) Time evolution of retrograde flow speed. (H) Total force acting on the substrate by the network with different $k_{20}$. Flow speed and total substrate force were proportional to $k_{20}$ at $k_{20} \leq 17$ s$^{-1}$ and reached a plateau at $k_{20} > 17$ s$^{-1}$.

**Table S1. List of parameters used for the lamellipodium model. Each parameter was tested prior to finding the reference case.** Some parameters were determined based on prior studies.

| **Symbol** | **Definition** | **Value** |
|---|---|---|
| $r_{0,\mathrm{A}}$ | Length of an actin segment | $1.4\times10^{-7}$ [m] |
| $r_{\mathrm{c,A}}$ | Diameter of an actin segment | $7.0\times10^{-9}$ [m] (82) |
| $\theta_{0,\mathrm{A}}$ | Bending angle formed by adjacent actin segments | 0 [rad] |
| $\kappa_{\mathrm{s,A}}$ | Extensional stiffness of F-actin | $1.69\times10^{-2}$ [N/m] |
| $\kappa_{\mathrm{b,A}}$ | Bending stiffness of F-actin | $2.64\times10^{-19}$ [N·m] (83) |
| $r_{0,\mathrm{ACP}}$ | Length of an ACP segment | $2.35\times10^{-8}$ [m] [81] |
| $r_{\mathrm{c,ACP}}$ | Diameter of an ACP segment | $1.0\times10^{-8}$ [m] |
| $\theta_{0,\mathrm{ACP}}$ | Bending angle formed by two ACP segments | 0 [rad] |
| $\kappa_{\mathrm{s,ACP}}$ | Extensional stiffness of ACP | $2.0\times10^{-3}$ [N/m] |
| $\kappa_{\mathrm{b,ACP}}$ | Bending stiffness of ACP | $1.04\times10^{-19}$ [N·m] |
| $r_{0,\mathrm{Arp2/3}}$ | Length of an Arp2/3 segment | $3.85\times10^{-8}$ [m] (84) |
| $r_{\mathrm{c,Arp2/3}}$ | Diameter of an Arp2/3 segment | $1.0\times10^{-8}$ [m] |
| $\theta_{0,\mathrm{Arp2/3c}}$ | Bending angle formed by two Arp2/3 segments | 0 [rad] |
| $\theta_{0,\mathrm{Arp2/3m}}$ | Bending angle between an Arp2/3 segment and a mother filament | 1.57 [rad] |
| $\theta_{0,\mathrm{Arp2/3d}}$ | Bending angle between an Arp2/3 segment and a daughter filament | 0.35 [rad] |
| $\theta_{0,\mathrm{Arp2/3f}}$ | Bending angle between mother and daughter filaments | 1.22 [rad] |
| $\phi_{0,\mathrm{Arp2/3}}$ | Torsion angle between mother and daughter filaments | 0 [rad] |
| $\kappa_{\mathrm{s,Arp2/3}}$ | Extensional stiffness of Arp2/3 arms | $2.0\times10^{-3}$ [N/m] |
| $\kappa_{\mathrm{b,Arp2/3c}}$ | Bending stiffness for $\theta_{\mathrm{Arp2/3c}}$ | $1.0\times10^{-19}$ [N·m] |
| $\kappa_{\mathrm{b,Arp2/3m}}$ | Bending stiffness for $\theta_{\mathrm{Arp2/3m}}$ | $1.0\times10^{-18}$ [N·m] |
| $\kappa_{\mathrm{b,Arp2/3d}}$ | Bending stiffness for $\theta_{\mathrm{Arp2/3d}}$ | $1.0\times10^{-18}$ [N·m] |
| $\kappa_{\mathrm{b,Arp2/3f}}$ | Bending stiffness for $\theta_{\mathrm{Arp2/3f}}$ | $1.0\times10^{-18}$ [N·m] |
| $\kappa_{\mathrm{t,Arp2/3}}$ | Torsional stiffness of Arp2/3 | $1.0\times10^{-18}$ [N·m] |
| $r_{0,\mathrm{MB}}$ | Length of a motor backbone segment | $4.2\times10^{-8}$ [m] |
| $r_{0,\mathrm{M1}}$ | Length of the transverse spring for a motor arm | $1.35\times10^{-8}$ [m] |
| $r_{0,\mathrm{M2}}$ | Length of the longitudinal spring for a motor arm | 0 [m] |
| $r_{\mathrm{c,M}}$ | Diameter of a motor arm | $1.0\times10^{-8}$ [m] |
| $\theta_{0,\mathrm{M}}$ | Bending angle formed by motor backbone segments | 0 [rad] |
| $\kappa_{\mathrm{s,MB}}$ | Extensional stiffness of a motor backbone | $1.69\times10^{-2}$ [N/m] |
| $\kappa_{\mathrm{s,M1}}$ | Extensional stiffness for the transverse spring for a motor arm | $1.0\times10^{-3}$ [N/m] |
| $\kappa_{\mathrm{s,M2}}$ | Extensional stiffness for the longitudinal spring for a motor arm | $1.0\times10^{-3}$ [N/m] |
| $\kappa_{\mathrm{b,M}}$ | Bending stiffness of a motor backbone | $5.07\times10^{-18}$ [N·m] |
| $\kappa_{\mathrm{r,A}}$ | Strength of repulsive force between F-actins | $1.69\times10^{-3}$ [N/m] |
| $N_{\mathrm{h}}$ | Number of heads represented by a motor arm | 4 |
| $N_{\mathrm{a}}$ | Number of arms per motor | 8 |
| $k_{20}$ | Unbinding rate of motor head | 17.14 [$\mathrm{s}^{-1}$]* |
| $k_{\mathrm{n,A}}$ | De novo nucleation rate of actin | $0.2\times10^{-6}$ [$\mu\mathrm{M}^{-1}\mathrm{s}^{-1}$] |
| $k_{\mathrm{b,A}}$ | Branch nucleation rate of actin | 20 [$\mu\mathrm{M}^{-1}\mathrm{s}^{-1}$] |
| $k_{+,\mathrm{A}}$ | Polymerization rate of actin at the barbed end | 12 [$\mu\mathrm{M}^{-1}\mathrm{s}^{-1}$]* |
| $k_{-,\mathrm{A}}$ | Depolymerization rate of actin at the pointed end | 6 [$\mathrm{s}^{-1}$]* |
| $k_{+,\mathrm{Arp2/3}}$ | Binding rate of Arp2/3 | 50 [$\mu\mathrm{M}^{-1}\mathrm{s}^{-1}$] |

| | | |
|---|---|---|
| $k_{+,ACP}$ | Binding rate of ACP | 100 [$\mu M^{-1}s^{-1}$] |
| $C_A$ | Actin concentration | 250 [μM] |
| $R_{Arp2/3}$ | Ratio of Arp2/3 concentration to $C_A$ | 0.01* |
| $R_{ACP}$ | Ratio of ACP concentration to $C_A$ | 0.04* |
| $R_M$ | Ratio of motor concentration to $C_A$ | 0.004* |
| $k_{0,sev}$ | Severing rate constant | $10^{-45}$ [$s^{-1}$]* |
| $\lambda_{sev}$ | Sensitivity to F-actin angle | 1.0 [deg] * |
| $A_{FA}$ | Size of focal adhesion region relative to domain size | 0.35* |
| $\Delta t$ | Time step | $1.15\times10^{-5}$ [s] |
| $\mu$ | Viscosity of surrounding medium | $8.6\times10^{-1}$ [kg/m·s] |
| $k_BT$ | Thermal energy | $4.142\times10^{-21}$ [J] |
| $\kappa_{s,sub}$ | Extensional stiffness of a substrate | $10^{-4}$ [N/m] |
| $\kappa_{b,sub}$ | Bending stiffness of a substrate | $2.4\times10^{-20}$ [Nm] |
| $r_{0,sub}$ | Length of a substrate chain | $5.0\times10^{-8}$ [m] |
| $\theta_{0,sub}$ | Bending angle formed by substrate chains | 1.05 [rad] |
| $\kappa_{s,C}$ | Extensional stiffness of clutches | $1.0\times10^{-3}$ [N/m] |
| $k^0_{u,C}$ | Zero-force unbinding rate constant of ACP (slip part) | 0.115 [$s^{-1}$] |
| $x_{u,C}$ | Sensitivity of ACP unbinding to applied force (slip part) | $1.04\times10^{-10}$ [m] |
